# Dynamic trajectories of connectome state transitions are heritable

**DOI:** 10.1101/2021.05.24.445378

**Authors:** Suhnyoung Jun, Thomas H. Alderson, Andre Altmann, Sepideh Sadaghiani

**Affiliations:** Beckman Institute for Advanced Science and Technology, University of Illinois at Urbana-Champaign, Urbana, Illinois, 618201; Psychology Department, University of Illinois at Urbana-Champaign, Urbana, Illinois 61801; Centre for Medical Image Computing (CMIC), Department of Medical Physics, University College London, London, UK; Neuroscience Program, University of Illinois at Urbana-Champaign, Urbana, Illinois 61801

**Author notes:** Correspondence: Sepideh Sadaghiani. **Author Contributions:** SJ and SS designed research; SJ performed research; SJ analyzed data; SJ and SS wrote the paper; and SS, THA and AA contributed analytic expertise, theoretical guidance, paper revisions, and informed interpretation of the results. **Competing Interest Statement:** The authors declare no competing interest.

**Keywords:** dynamic functional connectivity, heritability, variance component modeling, twin study, hidden Markov modelling.

## Abstract

The brain’s functional connectome is dynamic, constantly reconfiguring in an individual-specific manner. However, which characteristics of such reconfigurations are subject to genetic effects, and to what extent, is largely unknown. Here, we identified heritable dynamic features, quantified their heritability, and determined their association with cognitive phenotypes. In resting-state fMRI, we obtained multivariate features, each describing a temporal or spatial characteristic of connectome dynamics jointly over a set of connectome states. We found strong evidence for heritability of temporal features, particularly fractional occupancy (FO) and transition probability (TP), describing the trajectory of state transitions. Genetic effects explained a substantial proportion of phenotypic variance of these features (*h^2^*=.39, 95% CI= [.24,.54] for FO; *h^2^=*.43, 95% CI=[.29,.57] for TP). Moreover, these temporal phenotypes were associated with cognitive performance. Contrarily, we found no robust evidence for heritability of spatial features of the dynamic states (i.e., states’ Modularity and connectivity pattern). Genetic effects may therefore primarily contribute to how the connectome *transitions* across states, rather than the precise spatial instantiation of the states in individuals. In sum, genetic effects impact the duration spent in each connectivity configuration and the frequency of shifting between configurations, and such temporal features may act as endophenotypes for cognitive abilities.

## Introduction

Inter-individual variability in the time-averaged (static) functional connectivity architecture of the human brain is subject-specific and predictive of cognitive abilities (Finn et al. 2015; Jalbrzikowski et al. 2020; Ousdal et al. 2020; Rosenberg et al. 2016). With the increasing availability of large-sample fMRI datasets, significant genetic contributions to this *static* large-scale connectome architecture have been established. Specifically, individual edge-wise connections and networks of the functional connectome have been identified as heritable (Ge et al. 2017; Reineberg et al. 2020). Moreover, topological properties of the static connectome, such as the modular organization, have been shown to constitute heritable subject-specific traits linked to behavioral and cognitive characteristics (Liu, Kohn, and Fernández 2019; Sinclair et al. 2015).

However, the static architecture captures only part of the functionally significant properties of the connectome. In fact, the functional connectome as measured by fMRI exhibits flexible reconfigurations over the course of seconds to minutes (Bassett et al. 2011; Shine et al. 2019). These reconfigurations can be described as changes in connectivity strength between specific sets of brain region-pairs, forming recurrent connectome states. Such functional connectome states hold great significance as their time-varying (dynamic) characteristics have been linked to behavior and cognition (Eichenbaum et al. 2020; Vidaurre, Smith, and Woolrich 2017).

Because some of the same behavioral and cognitive features linked to connectome dynamics are also heritable (Han and Adolphs 2020), the possibility emerges that the behaviorally relevant connectome dynamics may themselves be heritable. A single prior study provides some support for this exciting possibility by demonstrating heritability of the ratio of time spent across two “meta-states” and their association with cognitive abilities (Vidaurre, Smith, and Woolrich 2017). However, beyond this single feature, it is unknown whether other properties of connectome dynamics are heritable. Further, the magnitude of heritability, i.e., % variance of the dynamic features explained by genetic effects, is unknown. We argue that future advances in genetics neuroimaging of connectome dynamics and its translational potential are contingent upon identifying which dynamic features are heritable and linked to behavior. Therefore, in the following we focus on a set of dynamic connectome features that have been most commonly reported as behaviorally relevant. This hypothesis-driven approach allows us to home in on a select subset among the numerous possible network features.

Particular *temporal* features of connectome dynamics, denoting durations and rates of states and their transitions, have been shown to be behaviorally relevant (Nomi et al. 2017). Specifically, the proportion of the total recording time spent in each connectome state (fractional occupancy, or FO) and the probability to transition between specific pairs of discrete states (transition probability, or TP) have been linked to behavior (Eichenbaum et al. 2020; Vidaurre, Smith, and Woolrich 2017). These temporal features are of particular interest, as they define the *trajectory* of the connectome through state space rather than the states themselves. In fact, the matrix of TP values between all possible state-pairs fully characterizes connectome transitions.

Beyond these temporal attributes of connectome reorganization, an often-reported observation is that dynamic changes in the *spatial* features of the connectome shape behavior. By spatial features, we refer to patterns of edge strength and ensuing topological characteristics of connectome states. In particular, behavioral outcomes have been linked to variability of functional connectivity (FC) between the default-mode network (DMN) and top-down control regions (Sadaghiani et al. 2015; Thompson et al. 2013). Most notably, FC dynamics between the DMN and the frontoparietal network (FPN) have been linked to cognitive flexibility (Douw et al. 2016; Hellyer et al. 2014; Vatansever et al. 2017). Spatial connectome patterns can further be quantified as global topological properties of the connectome’s graph. Modularity is the topological feature whose dynamics (Betzel et al. 2016) are most commonly associated with various behaviors (J. R. Cohen and D’Esposito 2016; Finc et al. 2017; Sadaghiani et al. 2015; Shine et al. 2016). Modularity quantifies the balance between segregation (prioritizing processing within specialized networks) and integration (combining specialized information from various networks) (Shine and Poldrack 2018). While modularity of the *static* connectome was found to be heritable (Sinclair et al. 2015), genetic contributions to the behaviorally-relevant Modularity dynamics across connectome states have not been assessed.

Based on the preceding evidence for behavioral relevance, we chose to investigate two dynamic temporal features, FO and TP, and two dynamic spatial features, time-varying FC between DMN and FPN (TV-FCDMN-FPN) and time-varying Modularity (TV-Modularity). We sought to answer (i) whether the hypothesis-driven temporal and spatial features of connectome dynamics are heritable; (ii) whether such heritability emerges from *multivariate* phenotypes jointly encompassing all connectome states or contrarily as phenotypes of individual states; (iii) how much of the phenotypic variance of the dynamic connectome features can be accounted for by genetic influence; and (iv) how much of the variance in heritable dynamic connectome features is associated with the individual variability in cognitive domains. To address these questions, we extracted discrete brain states from resting-state fMRI data acquired from the Human Connectome Project (Smith, Beckmann, et al. 2013), including monozygotic and dizygotic twin pairs, non-twin sibling pairs, and pairs of unrelated individuals, estimated their dynamic features, fitted quantitative genetic models to the features, and quantified their association with cognition.

## Results

Fig. 1 is a schematic representation of the overall approach and analysis subsections. Because neuroimaging data processing inevitably involves numerous decision points, we provide reasoning for our choices, and further include supplementary results for several alternative choices where appropriate (see *Supplementary Information*).

**Fig. 1.**
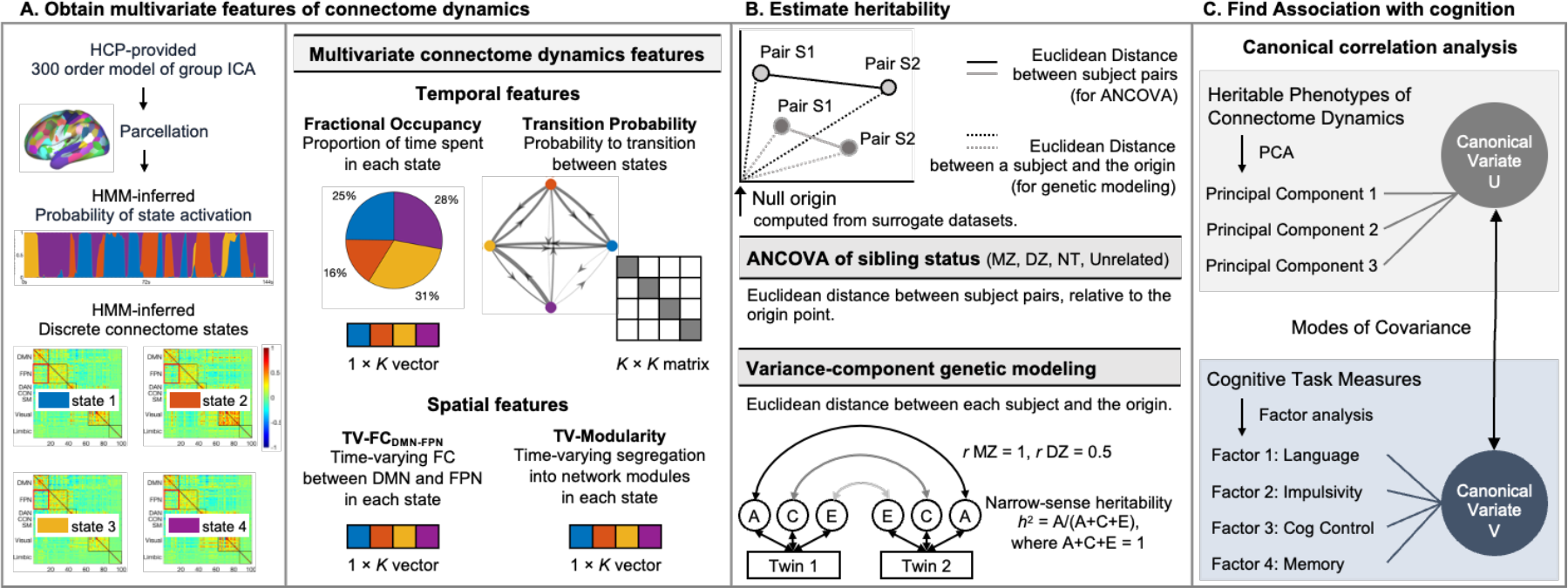
An overview of the analysis pipeline. We used minimally preprocessed resting state BOLD timeseries from 139 group-ICA-derived regions covering cortical and subcortical areas of the cerebrum as provided by the Human Connectome Project. [A] We used a hidden Markov model (HMM) to extract four discrete connectome states (or six states for replication) associated with a state time course for each subject indicating the probability of when each state is active. Multivariate temporal features of connectome dynamics were defined as the proportion of the recording time spent in each connectome state (fractional occupancy, or FO) and the probability matrix of transitioning between every pair of discrete states (transition probability, or TP). Multivariate spatial features included the time-varying functional connectivity between default mode and frontoparietal networks (TV-FCDMN-FPN) and time-varying global modularity (TV-Modularity). [B] We tested whether genetically more related subjects displayed greater similarity in their multivariate features than genetically less related subjects. First, for each feature of dimension *m*, we estimated a null model-derived origin point in the *m*-dimensional space. The position of each subject’s multi-dimensional feature value was estimated relative to this origin for genetic modeling (see below). Further, the similarity of this position between a given pair of subjects was quantified as Euclidean distance for ANCOVA analyses; a one-way ANCOVA of the factor sibling status with four levels (monozygotic twins (MZ), dizygotic twins (DZ), non-twin siblings (NT), and pairs of unrelated individuals) was performed on the distance value for each of the features. Secondly, we employed structural equation modeling (i.e., genetic variance component model) to quantify the genetic effects. Phenotypic variance of a trait was partitioned into additive genetic (denoted A), common environmental (denoted C) and unique environmental components (denoted E), with narrow-sense heritability (*h*^2^) quantified as the proportion of variance attributed to the genetic factor (A). Path A is dependent on the genetic similarity between twins. MZ twins are genetically identical (path denoted MZ = 1), whereas DZ twins based on the supposition of Mendelian inheritance share half of their genetic information (path denoted DZ = 0.5). [C] Finally, canonical correlation analysis (CCA) was used to find modes of population covariation between multivariate dynamic connectome features and cognition. CCA analysis was performed on dimension-reduced data including three multivariate dynamic connectome principal components and four cognitive factors

### I. Discrete connectome states have distinct spatial and temporal profiles

We extracted four discrete connectome states, i.e., whole-brain recurrent connectivity patterns, in a data-driven fashion using hidden Markov modeling (HMM) (Fig. 2A) (Vidaurre et al. 2016). HMMs require an *a priori* selection of the number of states (*K*), and prior HMM studies that have explored several *K*s have identified *K*s between 3 and 7 as optimal (Stevner et al. 2019; Vidaurre et al. 2016). Therefore, we chose *K of* 4 to fall within this range, and further assessed *K* of 6, reasoning that robust heritability effects would be evident regardless of the specific choice of K (within the optimal range). In the following, we show that robust heritability effects are observable in both parameter regimes. Here, we report outcomes for *K* = 4 and replicate results for *K* = 6 in *Supplementary Information* (*SI*).

**Fig. 2.**
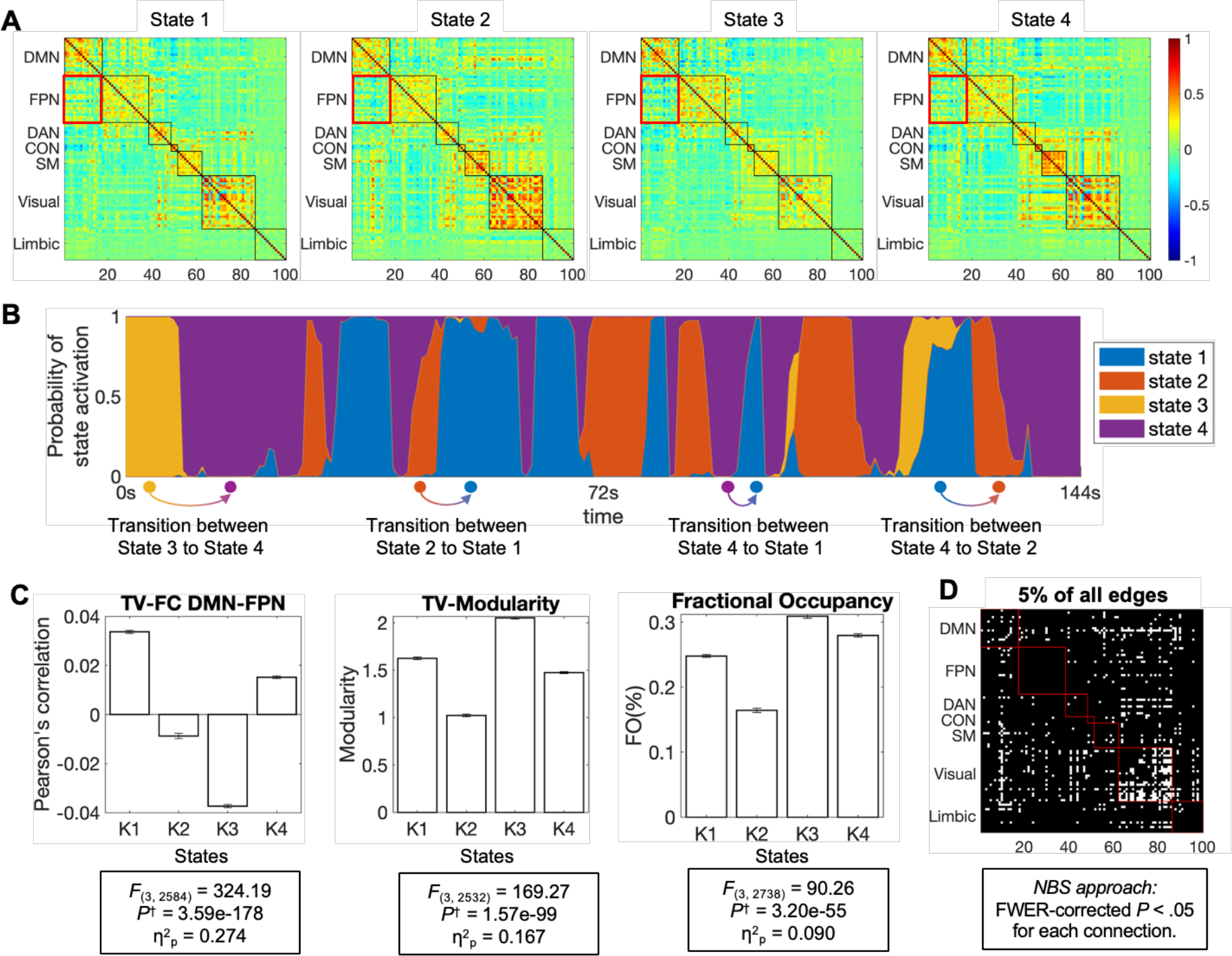
HMM states and state-dissociating features. (A) HMM estimates the connectome states that are common to all subjects. The states are represented by z-scored functional connectivity matrices. The rows and columns represent 139 regions organized according to canonical ICN membership (Yeo et al. 2011) including default mode network (DMN), frontoparietal network (FPN), dorsal attention network (DAN), salience/cingulo-opercular network (CON), sensory-motor network (SMN), visual network (VIS), and limbic network (Limbic; subcortical limbic regions omitted for visual succinctness). Red box indicates the state-specific functional connections between DMN and FPN (FCDMN-FPN) corresponding to an a priori selected spatial feature. (B) HMM estimates a specific (probabilistic) state time course for each subject indicating when each state is active. The states are characterized by their mean activation and functional connectivity matrix. An approximately two-minute section of the state time course is visualized for one subject exemplifying periods occupied by each state and transitions across states. (C) Bar plots of the mean time-varying FCDMN-FPN (TV-FCDMN-FPN), time-varying Modularity (TV-Modularity), and FO for each state (686 subjects). F values are reported for one-way ANCOVAs of the factor state for each variable. Strong differences across states are observed in all three measures. (*P*^†^: *P* values Bonferroni-corrected for three dependent variables. η^2^p: Partial Eta squared effect size). (D) A binary matrix showing a topological cluster of connections (edges) whose FC differed significantly across the four states. *F* values from an edge-wise ANCOVA of FC strength were threshold at 5% edge density followed by network-based statistics (NBS) to control for multiple comparisons (Zalesky, Fornito, and Bullmore 2010). Significant connections are widely distributed across networks, prominently including the DMN and VIS. For details, see *Fig. S5* and *SI Results IV*.

To assess whether the HMM-derived states possessed distinguishable network organization as expected, one-way ANCOVAs of factor state were conducted for each of the temporal and spatial dynamic features of interest, adjusting for age and head motion. Specifically, we tested the strength of FC between DMN and FPN (FCDMN-FPN), the degree of global modular organization as measured by Newman’s Modularity (Newman 2006), and the proportion of time spent in each state, or Fractional Occupancy (FO). Fig. 2B shows that the connectome states differed significantly with respect to FCDMN-FPN (*F*_(3, 2584)_ = 324.19, *P*^†^ = 3.57e-178, partial eta-squared (η^2^p) = .274, where *P*^†^ is the *P* value Bonferroni-corrected for three dependent variables), Modularity (*F*_(3, 2532)_ = 169.53, *P*^†^ *=* 1.13e-99, η^2^p = .167), and FO (*F*_(3, 2738)_ = 90.26, *P*^†^ = 3.20e-55, η^2^p = .090). An exploratory comparison of edge-wise FC strength further supported that FC differed across states in a spatially distributed set of connections (extending beyond FCDMN-FPN) (Fig. 2C; and *Fig. S5*). Regarding temporal characteristics, the distribution of FO over the states as well as the TP between states were not random (tested for the four- and six-state models against surrogate data, *SI Results I, Fig. S8*). These results confirm that the spontaneous connectivity time courses can be described as non-random sequences of four (or six) connectome states that differ from each other in terms of spatial organization, global topology, and proportion of occurrence.

### II. Multivariate temporal features of the dynamic connectome are heritable

Heritability was tested for *multivariate* features that concurrently characterize all states. To circumvent the statistical limitations imposed through multiple comparisons, the heritability analysis was confined to a selection of hypothesis-driven phenotypes. Specifically, the multivariate features included FO (1 × *K*), TP matrix (*K* × *K*), TV-FCDMN-FPN (1 × *K*), and TV-Modularity (1 × *K*).

We tested the supposition that genetically related subjects had more similar multivariate features than genetically less related subjects. The similarity of each of the multivariate connectome features between a given pair of subjects was quantified as Euclidean distance. Distance values entered a one-way ANCOVA of the factor sibling status with four levels, including monozygotic (MZ) twins, sex-matched dizygotic (DZ) twins, sex-matched non-twin (NT) siblings, and sex-matched pairs of unrelated individuals, adjusted for age and head motion. No subjects overlapped between groups.

We found that temporal features, describing the dynamic trajectory of connectome state transitions, are heritable (Fig. 3A). Specifically, genetically closer subject pairs have more similar FO (*F*_(3, 337)_ = 15.49, *P*^†^ = 7.32e-9, η^2^p = .121, where *P*^†^ is the *P* value Bonferroni-corrected for four dependent variables) and TP (*F*_(3, 337)_ = 12.00, *P*^†^ = 6.98e-7, η^2^p = .097) phenotypes, compared to less genetically-related pairs. This impact of sibling status on temporal features was of consistently large effect size (η^2^p ∼ .11) (J. Cohen 1988), regardless of global signal regression (GSR) and the chosen number of states (see *SI Results I*, *Fig. S2*). No effect of sibling status was observed in surrogate data lacking time-varying dynamics but with preserved static covariance structure; FO (*F*_(3, 337)_ = .66, *P*^†^ > 1.0, η^2^p = .006) and TP (*F*_(3, 337)_ = .82, *P*^†^ =.965, η^2^p = .007).

**Fig. 3.**
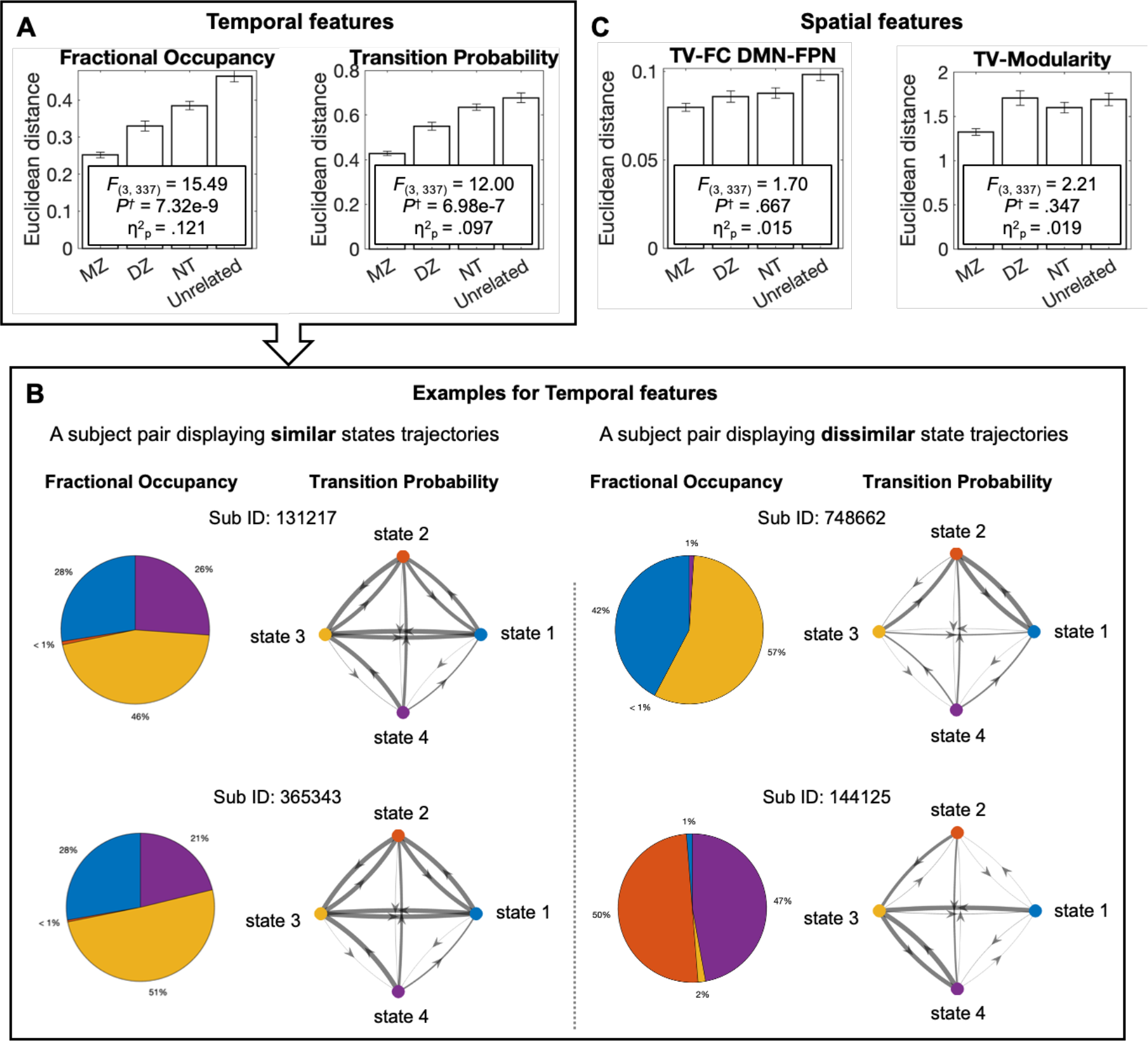
Heritability of multivariate dynamic connectome features. Heritability of each of the multivariate features was assessed by one-way ANCOVAs of the factor sibling status (four levels: monozygotic twins (MZ), dizygotic twins (DZ), non-twin siblings (NT), and pairs of unrelated individuals), adjusted for age and head motion. (A) Temporal connectome features, specifically the multivariate FO and TP phenotypes, were more similar among MZ twin pairs than DZ twins, followed by NT siblings and pairs of unrelated individuals. This influence of sibling status was statistically significant and stable, irrespective of global signal regression (GSR) and the chosen number of states (see *Fig. S2*). (B) To provide an intuition of the findings through examples, we illustrate FO and TP for the MZ subject pair with the highest similarity (*Left*), and for an unrelated subject pair with the lowest similarity in these features (*Right*). (C) In contrast to temporal features, spatial connectome features, including TV-FCDMN-FPN and TV-Modularity, did not show significant difference across sibling groups in the four-state model visualized here, and effect sizes remained small for alternative analysis choices (see *Fig. S3*). *P*^†^: *P* values Bonferroni-corrected for four dependent variables, η^2^p: Partial Eta squared effect size.

Contrary to the temporal features, we did not find robust support for heritability of the spatial features, which describe how connectome states are spatially instantiated in individuals. Specifically, outcomes of equivalent ANCOVAs for TV-FCDMN-FPN (*F*_(3, 337)_ = 1.70, *P*^†^ = .667, η^2^p = .015) and TV-Modularity (*F*_(3, 337)_ = 2.21, *P*^†^ = .347, η^2^p = .019) showed no impact of sibling status, and effect sizes remained small (η^2^p ∼ .02) irrespective of the chosen number of states and GSR (*SI Results I*, *Fig. S3*). This lack of heritability was confirmed by a subsequent variance-component genetic analysis (see **Results** III). Our sample size permitted detecting, at 80% power, effects of small size (η^2^ = .03, equivalent to *f* = .179, or larger). Therefore, if for spatial features heritability produced effects smaller than the detectable size, these effects would be of low practical impact.

To ensure that the lack of a robust outcome for spatial attributes was not driven by a narrow feature selection, we further performed exploratory ANCOVAs of sibling status for several additional features. Specifically, we assessed heritability of behaviorally relevant FC (Sadaghiani et al. 2015; Thompson et al. 2013) between DMN and the two other major top-down control networks (*SI Results III*, *Fig. S4*), as well as a data-driven selection of edges (*SI Results IV*, *Fig. S5*). Regardless of the chosen number of states and/or GSR, all exploratory spatial features showed no support for heritability or minimal effect size in this analysis. We performed additional Bayesian ANCOVAs to directly assess the probability of H0 (i.e. the null hypothesis that there is no effect of sibling status) against H1 (*Table S2*). Indeed, for TV-FCDMN-FPN a BF01 of 8.04 showed that the data are 8 times more likely to occur under H0 than under H1. For TV-Modularity, a BF01 of 2.15 provided anecdotal evidence in the direction of H0. Similarly, BF provided anecdotal to strong evidence for H0 for all other spatial features (cf. *Table S2*). These findings strongly contrast the observations for temporal features, where Bayes Factor of both FO and TP showed that the data are > 100 times more likely to occur under H1 than under H0 (BF01 < 10^-6^).

Importantly, the effect size of heritability was considerably larger for *multivariate* temporal features (Fig. 3) compared to the *individual* (i.e., state-by-state) components of the multivariate features (*Table S3*). Therefore, our findings demonstrate that the dynamic trajectory of connectome state transitions are heritable predominantly when considered as multivariate patterns, rather than as individual state-specific components.

### III. Genetic effects account for substantial variability in temporal connectome dynamics

We employed a structural equation modeling common in classical twin studies (Falconer 1990) to estimate how much of the phenotypic variance is explained by genetic variance, or heritability (*h*^2^). Traditionally, the model is fitted to univariate phenotypes. However, our above-described ANCOVAs suggest that temporal characteristics of connectome dynamics are inherited predominantly as *multivariate* phenotypes (Fig. 3A). Therefore, we adjusted the model to accommodate multivariate phenotypes by quantifying the similarity (or Euclidean distance) between multivariate features from each subject’s real data and a “null” point of origin (from dynamics-free surrogate data Fig. 1). To replicate our results independent of this novel approach, we provide results from an alternative approach for multivariate features (Ge et al. 2016), which however does not account for collinearities among the univariate components (*SI Results V* and *Table S4*).

A substantial portion of variance of temporal features was explained by additive genetic variance in the ACE model, adjusted for age, sex, and head motion (Table 1). The additive genetic effect (A, or narrow-sense heritability) of FO and TP was estimated as *h^2^* = .39 (95% confidence interval (CI): [.24, .54]) for FO and *h^2^ =*.43 (95% CI: [.29, .57]) for TP. Note that the impact of common environment (C) was estimated as zero for both FO and TP. For both features, the fitness of the nested models was not significantly better than the full ACE model. These outcomes indicate that genetics contribute substantially to the temporal features of connectome dynamics.

**Table 1.** Variance-component model parameter estimates of the dynamic connectome features.

| Phenotype | Model | $h^2$ | (95% CI) | A | (95% CI) | C | (95% CI) | E | (95% CI) | -2LL | AIC | $\chi^2$ | $\Delta df$ | p |
| --- | --- | --- | --- | --- | --- | --- | --- | --- | --- | --- | --- | --- | --- | --- |
| Temporal features of connectome dynamics |  |  |  |  |  |  |  |  |  |  |  |  |  |  |
| FO | ACE | .39 | (.24, .54) | .39 | (.24, .54) | .00 | (0.0, 0.0) | .61 | (.46, .76) | -528.84 | -516.84 |  |  |  |
|  | AE | .39 | (.20, .51) | .35 | (.20, .51) |  |  | .65 | (.49, .80) | -528.84 | -518.84 | 2.43e-9 | 1 | 1.0 |
|  | CE |  |  |  |  | .30 | (.17, .43) | .70 | (.57, .83) | -525.52 | -515.52 | 3.31 | 1 | .069 |
| TP | ACE | .43 | (.29, .57) | .43 | (.29, .57) | .00 | (0.0, 0.0) | .57 | (.43, .71) | -515.93 | -503.93 |  |  |  |
|  | AE | .43 | (.29, .57) | .43 | (.29, .57) |  |  | .57 | (.40, .71) | -515.93 | -505.93 | 2.05e-10 | 1 | 1.0 |
|  | CE |  |  |  |  | .35 | (.22, .47) | .65 | (.53, .78) | -512.78 | -502.78 | 3.15 | 1 | .076 |
| Spatial features of connectome dynamics |  |  |  |  |  |  |  |  |  |  |  |  |  |  |
| TV-FC<sub>DMN-FPN</sub> | ACE | .02 | (-.46, .49) | .02 | (-.46, .49) | .20 | (-.21, .61) | .78 | (.62, .94) | -1296.57 | -1284.57 |  |  |  |
|  | AE | .24 | (.08, .40) | .24 | (.08, .40) |  |  | .76 | (.60, .92) | -1295.94 | -1285.94 | .633 | 1 | .426 |
|  | CE |  |  |  |  | .21 | (.07, .35) | .79 | (.65, .92) | -1296.57 | -1286.57 | .003 | 1 | .957 |
| TV-Modularity | ACE | .05 | (-.54, .63) | .05 | (-.54, .63) | .07 | (-.36, .50) | .88 | (.66, 1.11) | 862.05 | 874.05 |  |  |  |
|  | AE | .14 | (-.06, .33) | .14 | (-.06, .33) |  |  | .86 | (.67, 1.06) | 862.15 | 872.15 | .095 | 1 | .758 |
|  | CE |  |  |  |  | .10 | (-.04, .24) | .90 | (.76, 1.04) | 862.08 | 872.08 | .026 | 1 | .872 |

All models were adjusted for age and head motion (FD). A, Additive genetic effect; C, common environmental effect; E, Unique/non-shared environment effect; -2LL, twice the negative log-likelihood; AIC, Akaike's information criterion; df, degrees of freedom; $\chi^2$ , chi square, $\Delta df$ , change in degree of freedom between the full model and the nested model; $p$ , $\chi^2$ test in model fitting. The AE and CE models are nested within the ACE model. Each nested model is compared with the fully saturated model. The fitness of models was tested based on a change in AIC (for a change of df of 1, the statistically significant change in $\chi^2$ is 3.84). $h^2$ , the narrow-sense heritability estimated as $\sigma^2_A/(\sigma^2_A + \sigma^2_C + \sigma^2_E)$ ; CI: Confidence Interval (lower bound, upper bound); FD, frame-wise displacement.

Consistent with the ANCOVA-based heritability findings (Fig. 3B and *Fig. S3*), the ACE model did not support genetic effects on either spatial feature (TV-FCDMN-FPN and TV-Modularity). The *h^2^* was estimated as .02 for TV-FCDMN-FPN and .05 for TV-Modularity, but with a wide 95% CI crossing zero: [-.46, .49] and [-54, .63], respectively.

### IV. Temporal phenotypes of the dynamic connectome are associated with cognition

Finally, for the heritable phenotypes of connectome dynamics, we assessed associations with cognitive measures using canonical correlation analysis (CCA) (Smith et al. 2015). CCA was performed on dimensionality-reduced data (three principal components for dynamic connectome phenotypes and four factors for cognitive measures). Among the three linear relationships (modes) of covariation, two modes were significant (against 10,000 permutations, *p* < 10^-4^, Fig. 4B and 4C) and robust after correcting for head motion. Post-hoc correlation between the modes and cognitive factors revealed the contribution of each factor to each mode. The most significant mode was defined by negative weights for the “Language” (*r* = -.20) and “Memory” (*r* = -.18) factors, followed by “Impulsivity” (*r* = -.14) and “Cognitive control” (*r* = -.12; Fig. 4D). The second significant mode was defined by a positive weight for “Cognitive control” (*r* = .07; Fig. 4E). Together, these findings suggest that temporal phenotypes of the dynamic connectome are linked to cognitive abilities.

**Fig. 4.**
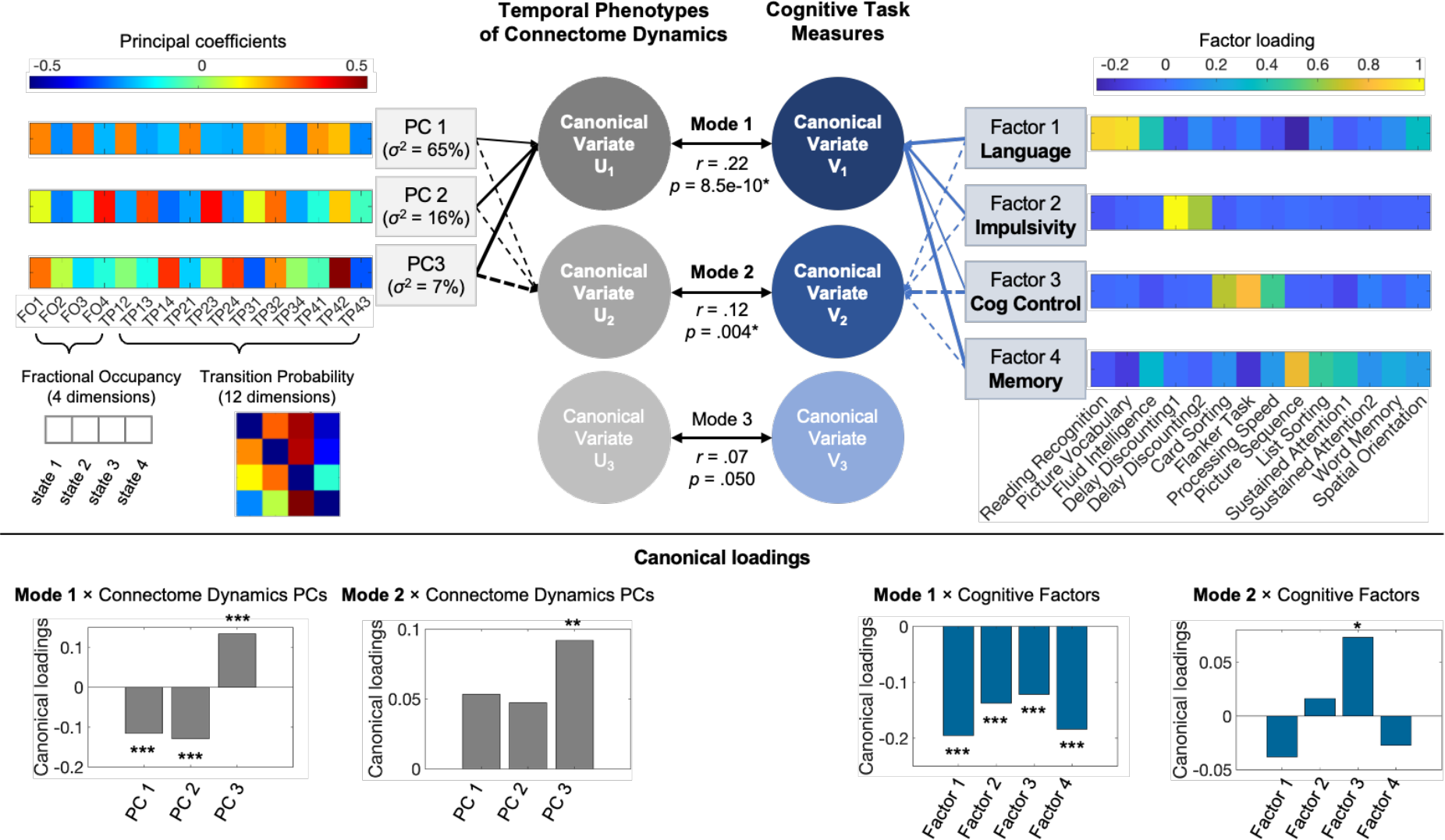
Temporal dynamic connectome features are related to cognition. Canonical correlation analysis (CCA) finds the maximum correlation between the linear combination between the two multi-dimensional canonical variates: U and V. Canonical variate U is defined as the three principal components (PCs), representing the temporal phenotypes of connectome dynamics. Canonical variate V is defined as four Factors, each representing the different domain of cognitive task measures. The principal coefficients matrix displays the weight that each component of the temporal phenotypes has on each of the PCs. The factor loading matrix displays the weight that each cognitive measure has on each of the Factors (see *Table S1* for details of cognitive task measures). CCA revealed two significant linear relationships (or modes) between the two canonical variates. The contribution of each PC or Factor to the given mode (as evaluated through post-hoc Pearson’s correlations with each mode) is illustrated by the thickness of the arrows linking PCs or Factors to each canonical variate. These contributions and their statistical significance are quantified by the bar plots in the lower panel. σ^2^: the variance explained by each principal component, Cog Control: cognitive control. * *p* < .05, ** *p* < .005, *** *p* < 5.0e-4.

## Discussion

The present interest in time-varying dynamics of the functional connectome is rooted in its impact on cognitive processes that are inherently dynamic (Kucyi et al. 2018), with implications for inter-individual differences in cognitive abilities (Eichenbaum et al. 2020; Nomi et al. 2017). Subject-specific features of connectome reconfigurations are both dynamic and multidimensional —they collectively incorporate patterns from multiple connectome states that evolve over time. However, the growing literature on heritability of connectome features has, for the most part, overlooked the connectome’s inherently dynamic and multi-dimensional character. Thus, we introduce a novel method providing comprehensive heritability assessments of a variety of multidimensional dynamic connectome features, and —for the first time<u>—</u> quantify the genetic effects on these multi-dimensional phenotypes (Table 1). This approach successfully demonstrates that heritability effect size is larger and more robust for multivariate dynamic features (I.e., FO, TP ;Fig. 2 and *Fig. S2*) than for state-wise scalar features (cf. example here; *Table S3*). . Taken together, our findings provide strong evidence for a substantial genetic effect on the dynamic trajectory of connectome state transitions that does not extend to the spatial and topological features of connectome states.

Previous studies based on the classical twin designs have found considerable genetic effects on multiple structural and *static* (i.e., time-averaged) functional connectome features of the human brain. For example, heritable features of the structural connectome include size (*h*^2^ = 23-60%) and topography (*h*^2^ = 12-19%) of ICNs (Anderson et al. 2021), and average regional controllability as derived from network control theory (*h*^2^ = 13-64%) (Lee et al. 2020). Regarding the functional connectome, heritability has been established for static FC within canonical neurocognitive ICNs in several studies (*h*^2^ = 13–36% (Adhikari et al. 2018); *h*^2^ = 45-80% (Ge et al. 2017); (*h*^2^ = 9-28% (Elliott et al. 2019)). Further, genetic effects have been reported for topological properties of the static functional connectome, such as global efficiency (*h*^2^ = 52-62%), mean clustering coefficient (*h*^2^ = 47-59%), small-worldness (*h*^2^ = 51-59%), and modularity (*h*^2^ = 38-59%) (Sinclair et al. 2015). The present multivariate approach represents a significant departure from previous studies which traditionally estimate heritability for univariate connectome features.

Beyond the above-described studies on structural and static functional connectomes, heritability and quantitative estimation of the genetic impact have been largely unknown for connectome dynamics. To our knowledge, only a single dynamic feature has undergone heritability analysis, specifically FO across a set of states and their binary meta-states (Vidaurre, Smith, and Woolrich 2017). The current study establishes heritability of a wider hypothesis-driven set of multivariate dynamic connectome phenotypes that encompasses both temporal and spatial dynamic features. The statistical effect sizes of heritability were consistently large across several different methodological choices for temporal characteristics of connectome state transitions (FO and TP). Further, we provided the first quantitative estimate of additive genetic effect on FO (*h^2^* = 39%) and TP (*h^2^* = 43%), using the classical twin design (Falconer 1990). This strong heritability is in the range reported above for structural and static FC investigations.

Interestingly, for both FO and TP, the effect of common environment was estimated to be zero (Table 1). The twin model assumes that environment affects MZ and DZ twins equally and, thereby, greater phenotypic similarity of MZ twins must be due to their greater genetic similarity. However, such biological assumptions can be violated as both intrauterine and postnatal environments can differ as a function of zygosity (Conley et al. 2013). Therefore, the impact of common environment on the phenotypic variance cannot be estimated with precision. Nonetheless, our estimated values suggest that the common environment is unlikely to have a sizable influence on these phenotypes.

Importantly, we showed that the chosen temporal characteristics of connectome state transitions are linked to cognitive abilities. Recently, the well-established association between cognition and structural (Lee et al. 2020) or static functional connectome features (Cole et al. 2013; Elliott et al. 2019; van den Heuvel and Sporns 2013) has been extended to functional connectome *dynamics* (Eichenbaum et al. 2020; Vidaurre, Smith, and Woolrich 2017). Our observations further corroborate the association between cognitive abilities and connectome dynamics. Although CCA-derived associations cannot reveal the mechanistic nature of this relationship (Eichenbaum et al. 2020), it has been suggested that connectome dynamics may facilitate cognitive processes that are inherently dynamic in nature (J. R. Cohen 2018). Notably, the association with dynamic *temporal* features, specifically, suggest that subject-specific *trajectories* across connectome states are of importance to cognitive processes, most notably language, memory, and cognitive control. Importantly, our observations further suggest genetic contributions to connectome dynamics that impact cognition. A potential driving force of the heritability of such temporal aspects of connectome dynamics could be variability in (e.g., receptor) genes of modulatory neurotransmitter systems. This possibility is in line with a leading theory suggesting that ascending neuromodulatory input may serve as a primary drive behind connectome dynamics and cognition (Shine et al. 2019; 2019). Therefore, the link between dynamic trajectories of the time-varying connectome and cognitive abilities may suggest that temporal phenotypes are potential endophenotypes for cognitive abilities and therefore suitable candidates for genetic association studies.

In contrast to the temporal features, spatial characteristics of connectome states (TV-FCDMN-FPN and TV-Modularity) displayed a lack of heritability or consistently small effect sizes (η^2^p ∼ .02; Fig. 3B and *Fig. S3*). Beyond our hypothesis-driven spatial features, we expanded our search for heritable spatial features to include TV-FCDMN-DAN, TV-FCDMN-CON, and data-driven clusters of connections. These exploratory spatial features likewise showed no evidence for heritability or only marginal effect sizes across different methodological choices (See *SI Results III* and *IV*). In principle, this null result may be driven by limited signal-to-noise ratio, limited statistical power, or other factors. However, the weight of evidence suggests that genetic effects primarily contribute to how the connectome *transitions* across different states, rather than the precise way in which the states are spatially instantiated in individuals.

Our study is subject to several limitations and methodological considerations. In principle, one might conceive of a scenario where genetic effects impact a mental process or a different neurobiological process that in turn affects connectome dynamics. In such a hypothetical case, we believe that our results would still be of basic and translational value as they would demonstrate that genetic effects contribute --via an indirect route-- to inter-individual differences in connectome dynamics. Another consideration is that the available sample size is relatively small for a heritability study. Despite this limitation, the confidence intervals indicate that the sample size was sufficient to establish the genetic effect on FO and TP with high confidence. Another limitation is that other spatial features that may be heritable may have been missed in this study. An exploratory study with a more exhaustive set of features (and respective multiple comparisons corrections) may be sub-optimal given the limited sample size. Thus, a hypothesis-driven set of dynamic connectome features based on previously established behavioral relevance was indicated and adopted here. Additionally, we reported the absence of heritability for FC between DMN and the two other major task-positive networks, CON and DAN (Sadaghiani et al. 2015; Thompson et al. 2013), as well as for a fully data-driven selection of graph edges. Larger sample sizes may provide insights into heritability of other spatial dynamic features in the future; our results provide a starting point for explorations of these larger feature sets.

In conclusion, our findings establish that transitions between whole-brain connectome states and the proportion of time spent in each state are heritable and subject to substantial genetic influence. These results also suggest a likely non-genetic origin for inter-individual differences in the spatial layout of connectome states. This evidence adds to previous findings linking heritable temporal dynamics of connectome states and cognition (Vidaurre, Smith, and Woolrich 2017; Eichenbaum et al. 2020) and identifies TP and FO in the resting human brain as potential endophenotypes for cognitive abilities. As such, these features may inform investigations into specific, functionally relevant genetic polymorphisms and translate to efficient connectome-based biomarkers.

## Materials and Methods

### Neuroimaging and Behavior Dataset

We used the Washington University-University of Minnesota (WU-Minn) consortium of the Human Connectome Project (HCP) S1200 release (Van Essen et al. 2013). Participants were recruited, and informed consent was acquired by the WU-Minn HCP consortium according to procedures approved by the Washington University IRB (Glasser et al. 2013). For details of the HCP data collection protocol and cognitive measures, see (Smith, Beckmann, et al. 2013; Van Essen et al. 2013) and (Barch et al. 2013), respectively.

From the pool of all 1003 healthy adult subjects (aged 22-37 y, 534 females) with four complete resting-state fMRI runs (4800 total timepoints) we investigated 120 monozygotic (MZ) twin pairs, 65 sex-matched dizygotic (DZ) twin pairs, 96 sex-matched non-twin (NT) sibling pairs, and 62 pairs of sex-matched unrelated individuals. Note that all pairs are uniquely defined so that none of the subjects overlap between groups to avoid dependencies across members of families with > 2 subjects. All 1003 subjects entered HMM estimation of discrete connectome states, while 686 subjects (as described in the pairs above) entered heritability analysis. We included all 14 cognitive measures, which are summary scores for either a cognitive task or a questionnaire, under the cognition domain provided by the HCP (see *Table S1* for more detailed description for each variable, and *Fig. S7A* for their phenotypic correlation structure). The measures were z-score normalized to zero mean and unit variance. Of 1003 subjects, 997 subjects had complete data for the 14 cognitive variables measuring cognitive performance and entered canonical correlation analysis (CCA).

### Neuroimaging Data Preprocessing

All imaging data were acquired on a customized Siemens 3 T Skyra at Washington University in St. Louis using a multi-band sequence. Each 15-minute resting-state fMRI run was minimally preprocessed (Glasser et al. 2013) using tools from FSL (Jenkinson et al. 2012) and Freesurfer (Fischl 2012), and had artifacts removed using ICA+FIX (Griffanti et al. 2014; Salimi-Khorshidi et al. 2014). Preprocessing following the pipeline of Smith et al. (Smith, Beckmann, et al. 2013; Smith, Vidaurre, et al. 2013). Inter-subject registration of cerebral cortex was carried out using areal-feature-based alignment and the Multimodal Surface Matching algorithm (‘MSMAll’) (Glasser et al. 2016; Robinson et al. 2014).

### Parcellation

The HCP team has previously parcellated the neuroimaging data with ICA in FSL using various model orders of 25, 50, 100, 200, and 300 independent components (ICs). Specifically, for group-ICA, each dataset was temporally demeaned and variance normalized (Beckmann and Smith 2004). HCP provides averaged BOLD time-series for regions of these group-ICA-based parcellations, both with and without global signal regression.

Among the available parcellations, the 300 model order ICA was chosen for this study due to its following advantages: (i) ICs in this parcellation better separate the *individual* brain areas of the intrinsic connectivity networks (ICNs) than the lower model orders; (ii) in this parcellation those ICs with multiple brain regions spatially overlaps with a single ICN, whereas in lower model orders, ICs may overlap with multiple ICNs, making it difficult to consider the BOLD time-series extracted from a given IC as representing a single ICN.

However, the 300-model order ICA resulted in additional extreme parcellation of the cerebellum and brainstem; 161 out of 300 ICs were identified as either cerebellar or brainstem using both Automated Anatomical Labeling (AAL) atlas-based masks and visual inspection. Including these regions would underestimate the contribution of cortical and subcortical ICNs to the dynamic reconfiguration of the connectome. Therefore, we focused our investigation on the ICNs by excluding ICs which fall into the cerebellar/brainstem regions, resulting in a total of 139 out of 300 ICs.

We computed the spatial overlap between each of the 139 ICs and Yeo’s canonical neurocognitive ICNs (Yeo et al. 2011) to identify the functional network represented by each IC. Yeo’s 7 ICNs include visual network (VIS), sensory-motor network (SMN), dorsal attention network (DAN), salience/cingulo-opercular network (CON called Ventral Attention by Yeo et al.), limbic network (Limbic), frontoparietal network (FPN), and default mode network (DMN). While Yeo’s 7 ICNs are limited to cortical surface maps, the Limbic system is well-known to include core subcortical regions. To add these regions to our Limbic network, we selected the respective ICs that overlapped with an additional subcortical mask including the bilateral hippocampus, caudate, putamen, pallidum, and thalamus regions from the AAL atlas (see *Fig. S1* for the ICs grouped into the 7 ICNs).

### Hidden Markov modeling

We applied a hidden Markov model (HMM) to the minimally preprocessed region-wise BOLD time-series, resulting in *K* discrete states of whole-brain connectivity, each associated with state-specific time-course. The HMM assumes that time series data can be described using a finite sequence of a hidden number of states. Each state and its associated time series comprise a unique connectivity pattern that temporally re-occurs over time. Using the HMM-MAR (multivariate autoregressive) toolbox (Vidaurre et al. 2016), discrete connectome states were inferred from region-wise BOLD time-series temporally concatenated across all subjects. Whereas the states are estimated at the group-level, each individual has a subject-specific state time course, indicating when a given state is active. For a given K, the best fitting model corresponding to the one with the lowest free energy was selected across five runs (Quinn et al. 2018; Vidaurre et al. 2016; 2018).

HMMs require an a priori selection of the number of states, *K*. Generally, the objective is not to establish a ‘correct’ number of states but to strike a balance between model complexity and model fit and to identify a number that describes the dataset at a useful granularity (Quinn et al. 2018). Prior HMM studies that have directly compared several *K*s have identified *K*s between 3 and 7 as optimal (Stevner et al. 2019; Vidaurre et al. 2016). Therefore, we chose *K*=4 to fall within this range, and further assessed *K*=6, reasoning that robust heritability effects would be evident regardless of the specific choice of *K* (within the optimal range).

### Null model

We demonstrated that the dynamic trajectory of connectome state transitions is not random by generating 50 simulated state time courses of the same length as the original empirical state time courses (Vidaurre et al. 2016). Note that fifty is a robust number compared to previous work (e.g., Vidaurre et al. (Vidaurre, Smith, and Woolrich 2017) applied four simulations). The surrogate data, simulated using the *simhmmmar* function provided by HMM-MAR toolbox, preserve the static covariance structure of the original data but destroy the precise temporal ordering of states. An HMM inference with *K* = 4 and *K* = 6, respectively, was run on each of 50 surrogate datasets and the dynamic connectome features (FO, TP, etc.) at the group- and subject-level were recomputed. We confirmed that the non-random distribution of features over states observed in the original dataset was absent in the surrogate dataset (*Fig. S8)*.

### Deriving multivariate dynamic connectome features

For each subject and HMM-derived state, we calculated FO (the cumulative total time spent in a given state), FCDMN-FPN (the mean connectivity across all regions of DMN with those of FPN), and Newman’s Modularity (quantifying the level of segregation of each connectome state into modules (Newman 2006), which were configured to use canonical ICNs (Yeo et al. 2011) as the modular partition (Brain Connectivity Toolbox (Rubinov and Sporns 2010)). Further, for each subject, we estimated the transition probabilities across all state-pairs (TP matrix). All estimated variables were used as *multivariate* features that concurrently characterize all states. Therefore, the features included FO (1 × *K*), TP matrix (*K* × *K*), TV-FCDMN-FPN (1 × *K*), and TV-Modularity (1 × *K*).

### Similarity estimation and heritability testing

The multidimensional space for each feature was created by setting the origin point to the average of the given multivariate feature from the 50 surrogate datasets (see null model above). Within the multidimensional space constructed for each of the multivariate features, we estimated the similarity of each multivariate feature based on the Euclidean distance between a given pair of subjects (Colclough et al. 2017). Crucially, this similarity estimation approach preserved the positional relationship between elements in each multivariate variable. We tested if the similaritym of each of the multivariate features was dependent on the similarity of the genetic makeup between a given subject pair. Specifically, we used a one-way ANCOVA of the factor sibling status with four levels (i.e., MZ twins, DZ twins, NT siblings, and pairs of unrelated individuals) on the similarity of each of the multivariate features, adjusting for age and head motion similarity between subject pairs. Head motion was quantified as framewise displacement (FD) (Power et al. 2012), once in terms of the multivariate FD pattern across states (used for all analyses reported in the main manuscript, cf. *SI Results II* for details), and alternatively as FD across the total scan duration (reported in *SI Results II*).

Further, we examined if heritability of features was driven by the multivariate pattern or by *individual* (i.e., state-by-state) components of the multivariate features. Specifically, we estimated the similarity of each state-specific component of the FO, TV-FCDMN-FPN and TV-Modularity between a given pair of subjects. Likewise, the similarity of each state-pair component of the TP matrix was estimated. We used two-way ANCOVAs to examine the effect of sibling status and connectome state on the similarity of individual components of the multivariate features.

### Quantification of heritability

Heritability (*h^2^*) is the proportion of variance of a phenotype explained by genetic variance. We employed a structural equation modeling commonly used in classical twin studies to estimate heritability applied in twin studies. With multiple biological assumptions (Keller and Coventry 2005), including that environment affects MZ and DZ twins equally, the structural equations modeling partitions the phenotypic variance into the three components (A, C, and E) using maximum likelihood methods: additive genetic variance (A; resulting from additive effects of alleles at each contributing locus), common environmental variance (C; resulting from common environmental effects for both members of a twin pair), and random environmental variance (E; resulting from non-shared environmental) (Yashin and Iachine 1995).

The ACE model finds the correlation of univariate phenotypes between twin pairs and requires each subject to have a singular value for each phenotype. To accommodate the multivariate phenotypes to be fitted into the model, we defined an origin point from which the Euclidean distance of each multivariate phenotype will be estimated for each subject (Fig. 1).

This method was implemented in the *R* package *mets* (http://cran.r-project.org/web/packages/mets/index.html), adjusting for age and head motion. Nested models, specifically AE and CE, were fitted by dropping C or A, respectively. Statistical significance of nested models was assessed by a likelihood ratio test and the fitness of models was tested using the Akaike’s Information Criterion (AIC) (Akaike 1987).

### Relating dynamic trajectories of states to cognitive performance

Canonical correlation analysis (CCA) is a multivariate technique that can identify and measure linear relations between two sets of variables (Hotelling 1936). The CCA uses the Rao’s approximation approach to test the null hypothesis that each linear relationship (or “mode”) between the U and V canonical variates is zero. As a result, the significance of each mode is estimated. As previously applied to the static connectome and HMM-derived states, we trained CCA on the dimensionality-reduced temporal phenotypes of the connectome dynamics (U canonical variate matrix) and cognitive measures (V canonical variate matrix) (Eichenbaum et al. 2020; Smith et al. 2015; Vidaurre, Smith, and Woolrich 2017; Tibon et al. 2021).

Specifically, we reduced the dimensionality of the U and V canonical variates as follows. First, we applied a principal component analysis (PCA) to the U variate consisted of 16 temporal phenotypes of connectome dynamics (i.e., 4 dimensions from 1 × 4 FO and 12 dimensions from the off-diagonals of 4 × 4 TP matrix). We retained principal components (PCs) that had eigenvalues > 1 and identified three principal components (PCs) together explaining about 88.27% of the total variance in the temporal phenotypes of dynamic connectome (*Fig. S6*).

The dimensionality of V matrix consisted of 14 cognitive measures was reduced by following the dimensionality-reduction protocol used in the previous study that used HCP provided behavioral measures to investigate their heritability (Han and Adolphs 2020). After identifying the four PCs with eigenvalues > 1 (*Fig. S7B*), we performed a factor analysis to cluster the cognitive measures into four factors based on maximum likelihood estimates of the factor loadings (*Fig. S7C*). Factors were rotated using Promax oblique rotation because there was no evidence that the factors were orthogonal (*Fig. S7A*). We also computed factor scores using both regression and Bartlett methods for reliability (since factor scores are indeterminate). These two methods produced two sets of very similar factor scores for the same four factors (see correlation structure between all 8 factor scores in *Fig. S7D*). We then used the four factor scores from the regression method as the Y canonical variate. Based on the factor loading matrix showing the variance explained by each cognitive measure on a given factor (*Fig. S7C*) (Han and Adolphs 2020), we cautiously (and inevitably subjectively) interpret the extracted four factors as follows: Factor 1: “Language” (indicated by high positive loadings on language-related cognitive tasks); factor 2: “Impulsivity/self-regulation” (indicated by high positive loadings on delay discounting tasks); factor 3: “Cognitive control” (indicated by high positive loadings on fluid ability measuring tasks); and factor 4: “Memory” (indicated by high positive loadings on episodic and working memory tests).

To assess the statistical significance of the discovered modes of covariation, we calculated 10,000 permutations of the rows of U relative to V, respecting the within-participant structure of the data, and recalculated the CCA mode for each permutation in order to build a distribution of canonical variate pair correlation values (Smith et al. 2015). By comparing the outcome from the CCA of the true data to the shuffled data, we found that each mode of covariation discovered with the true data was highly significant (*p* < 1/10,000). The contribution of each factor to the given mode was evaluated with post-hoc correlations between the modes and the cognitive factors.

## Acknowledgments

Data were provided by the Human Connectome Project, WU-Minn Consortium (Principal Investigators: David Van Essen and Kamil Ugurbil; 1U54MH091657) funded by the 16 NIH Institutes and Centers that support the NIH Blueprint for Neuroscience Research; and by the McDonnell Center for Systems Neuroscience at Washington University. We would like to thank Jaime Derringer for her insight and interpretation of quantitative genetic modeling results and Lili Sahakyan for her insightful feedback on the manuscript. AA holds an MRC eMedLab Medical Bioinformatics Career Development Fellowship. This work was partly supported by the Medical Research Council (grant number MR/L016311/1). This work was supported by the National Institute for Mental Health (1R01MH116226 to Sepideh Sadaghiani).

## Supplementary Information

### Supplementary Results

#### SI Results I. Independence of heritability from parameter choices

To further support the robustness of the heritability of multivariate temporal features of dynamic functional connectome, we tested whether the heritability is independent of parameter choices: (i) global signal regression (GSR) and (ii) the chosen number of states. We repeated the one-way ANCOVAs of the factor sibling status with the four levels (MZ, DZ, NT, and Unrelated) on the connectivity data of the same subjects with GSR performed and with a different number of states (*K* of 6), adjusting for age and head motion.

*Fig. S2* shows that in all conditions, temporal dynamic features (i.e., TP and FO) are more similar among MZ twin pairs, followed by DZ twins and then NT siblings, and all the above showing higher similarity than pairs of unrelated individuals. This outcome demonstrates that heritability of the dynamic connectome features is robust and independent of GSR and the number of states *K*.

In contrast to these temporal features, *Fig. S3* shows that the heritability of the spatial dynamic features TV-FCDMN-FPN and TV-Modularity is dependent on the chosen number of states and GSR and of small effect size. Moreover, we found inconsistencies for the spatial features that may arise from noise in the data. Specifically, we observed that DZ twins have less similarity than NT siblings in some conditions. This pattern is inconsistent with expectations, given that DZ twins are subject to more strongly shared environmental factors than NT siblings.

#### SI Results II. Accounting for head motion

It is important to control for head motion in investigations of connectivity and its dynamics. In the context of our multivariate features of interest, which span several connectome states, a logical approach is to assess head motion in a likewise multivariate manner, thus accounting for the *pattern* of head motion over the different states. To this end, we used the framewise displacement (FD) (Power et al. 2012) computed from head motion parameters provided by the HCP and averaged them over all occurrences of each state. We then estimated the similarity of the multivariate FD vector (1 × *K* states) as the Euclidean distance between a given pair of subjects, paralleling the approach used for the dynamic features of interest. We did not find evidence for the heritability of multivariate FD (*K* = 4: *F*_(3, 338)_ = 1.32, *P* = 0.27, η^2^p = .012; K = 6: *F*_(3, 338)_ = .597, *P* = 0.617, η^2^p = .005), after adjusting for age. Nevertheless, for all heritability analyses of dynamic connectome features reported in the main manuscript, we controlled for multivariate FD as a covariate to account for any potential effect of head motion.

Additionally, since head motion across the *entire* scan time has previously been found to be heritable in the HCP data (Hodgson et al. 2017), we repeated our core heritability analyses (corresponding to Fig. 3 and Table 1 in the main text) using the FD obtained across the entire scan time. In line with the main report (Fig. S3A), we found consistently significant and large effects of the factor sibling status on temporal features. Specifically, genetically closer subject pairs have more similar FO (*F*_(3, 337)_ = 12.80, *P*^†^ = 2.44e-7, η^2^p = .102, where *P*^†^ is the *P* value Bonferroni-corrected for four dependent variables) and TP (*F*_(3, 337)_ = 10.18, *P*^†^ = 7.81e-6, η^2^p = .083) phenotypes, compared to less genetically-related pairs. In line with the main report (Fig. S3B), we did not find robust support for heritability of the spatial features. Specifically, outcomes of equivalent ANCOVAs for TV-FCDMN-FPN (*F*_(3, 337)_ = 1.50, *P*^†^ = .855, η^2^p = .013) and TV-Modularity (*F*_(3, 337)_ = 2.12,*P*^†^ = .392, η^2^p = .019) showed no impact of sibling status. Further, consistent with the main subsequent variance-component genetic analysis (Table 1), the ACE model estimated the additive genetic effect of FO and TP as *h^2^* = .34 (95% CI: [.18, .49]) for FO and *h^2^ =*.38 (95% CI: [-.13, .88]) for TP.

#### SI Results III. Exploring heritability of DMN connectivity to other networks

To extend our understanding of spatial features, additional to the TV-FCDMN-FPN and TV-Modularity, we performed exploratory ANCOVAs to assess the heritability of FC between DMN and the two other major task-positive networks, cingulo-opercular (CON) and dorsal-attention (DAN) networks (Sadaghiani et al. 2015; Thompson et al. 2013). Repeating the process in the *SI Results I*, we applied one-way ANCOVAs of the factor sibling status with the four levels (MZ twins, DZ twins, NT siblings, and pairs of unrelated individuals) on these additional features. *Fig. S4* shows that the effect size of heritability of both TV-FCDMN-CON and TV-FCDMN-DAN are small, regardless of non-GSR/GSR and the chosen number of *K*. Additionally, the ACE model showed that no heritability is supported for both TV-FCDMN-CON and TV-FCDMN-DAN.

#### SI Results IV. Exploring heritability with network-based statistics

Further expanding our selection of features in a data-driven manner, we adopted the network-based statistics (NBS) approach to identify topological clusters that are significantly different across discrete connectome states in a data-driven manner. NBS is a non-parametric method that deals with the multiple comparisons problem on a graph by identifying the largest connected sub-component (edge cluster) in topological space while controlling the family-wise error rate (FWER) (Zalesky, Fornito, and Bullmore 2010).

First, we performed mass univariate *F*-testing across four connectome states, independently across every connection within the connectome. Importantly, *F* tests were performed on absolute values for all connections to focus on the strength of connections, regardless of their connectivity direction. Each connection is endowed with a single test statistic. Then, the test statistic is thresholded at an arbitrary value to obtain a set of supra-threshold connections, then identify topological clusters among the set of supra-threshold connections. Since the choice of threshold is arbitrary, we chose a range of thresholds that allow 1% to 5% of the entire connections to survive. Most notably, because NBS clusters are defined in topological space, not in physical space, a connected path can in principle be found between any two nodes. Finally, FWER-corrected *P*-value (*p* < .05) for each component is tested using permutation testing (*n* = 5000). The permutation constructs an empirical null distribution of the largest connected cluster size by randomly rearranging the correspondence between data points and their labels under the null hypothesis without affecting the test statistic. We measured the size of a component as the total number of connections or extent of that component, as it is appropriate for detecting relatively weak effects that extend to encompass many connections.

After identifying the clusters for each threshold in four- and six-state models, respectively, we performed one-way ANCOVAs of the factor sibling status to assess the heritability of the absolute connectivity strength averaged over all supra-threshold edges. *Fig. S5* shows that the heritability is not significant across the selected thresholds in both *K*s of 4 and 6 models.

#### SI Results V. Alternative multivariate genetic variance component model

One of the novelties of our work is that we provided a novel approach to adjust the common structural equation model traditionally used in classical twin studies of univariate phenotypes (Falconer 1990) to fit multivariate phenotypes. This adjustment was made available by using the similarity (or Euclidean distance) between subject’s multivariate feature and a given origin point (**Materials and Methods** – Quantification of heritability; Fig. 1). To provide further support for the validity of this adaptation, we applied an alternative model (Ge et al. 2016) to estimate the heritability of multivariate features. This alternative weighted sum approach computes heritability of the multivariate phenotype as the cumulative heritability of individual components (cf. *Table S4* for formula). However, it should be noted that this cumulative approach will overestimate heritability for highly collinear phenotype features (since the shared variance explained by collinear features enters the sum multiple times), and as such is not optimal for FO and TP.

Despite this conceptual difference to our multivariate model, the cumulative heritability model resulted in comparable outcomes. The sum of weighted heritability of FO was calculated as .48 and of TP as .38, confirming strong heritability (*Table S4)*. These values are reasonably close to (within the confidence intervals of) those estimated in our multivariate approach. The converging results support the validity of our multidimensional space-based approach, including its origin point determined from surrogate data.

## Supplementary Figures

**Fig. S1.**
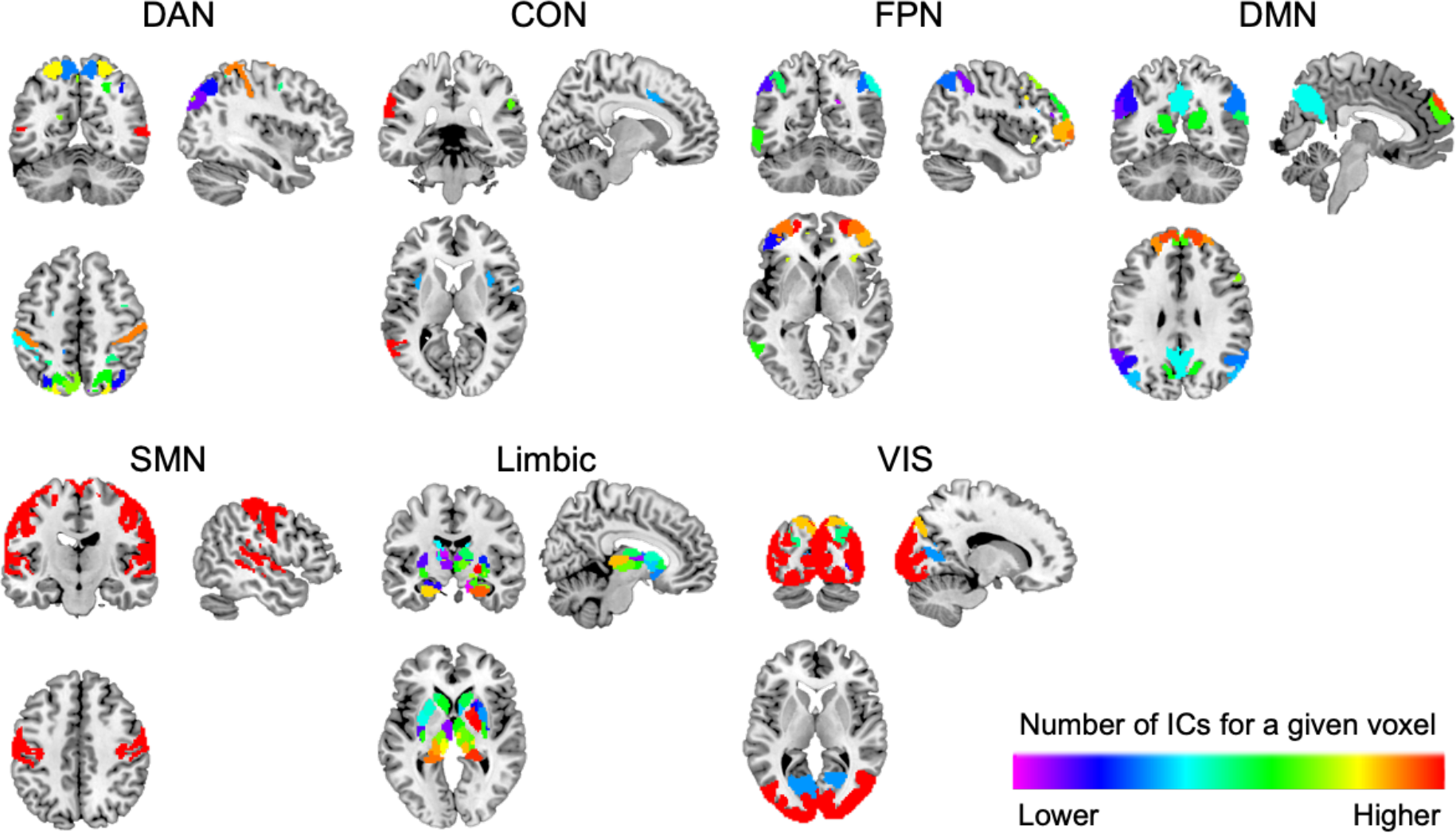
Cortical spatial maps of the selected 139 independent components of the 300-parcel model provided by HCP. Cortical spatial maps representing the selected 139 independent components (ICs) have been grouped into Yeo’s seven functional domains: dorsal attention network (DAN), salience/cingulo-opercular network (CON), frontoparietal network (FPN), default mode network (DMN), sensorimotor network (SMN), Limbic network, and visual network (VIS). Details of the parcellation and IC selection criteria are described in the **Materials and Methods** – Parcellation. The color bar indicates the number of ICs (out of 137 ICs) corresponding to the given voxel. The hotter the color is, the more ICs overlap with a given ICN mask.

**Fig. S2.**
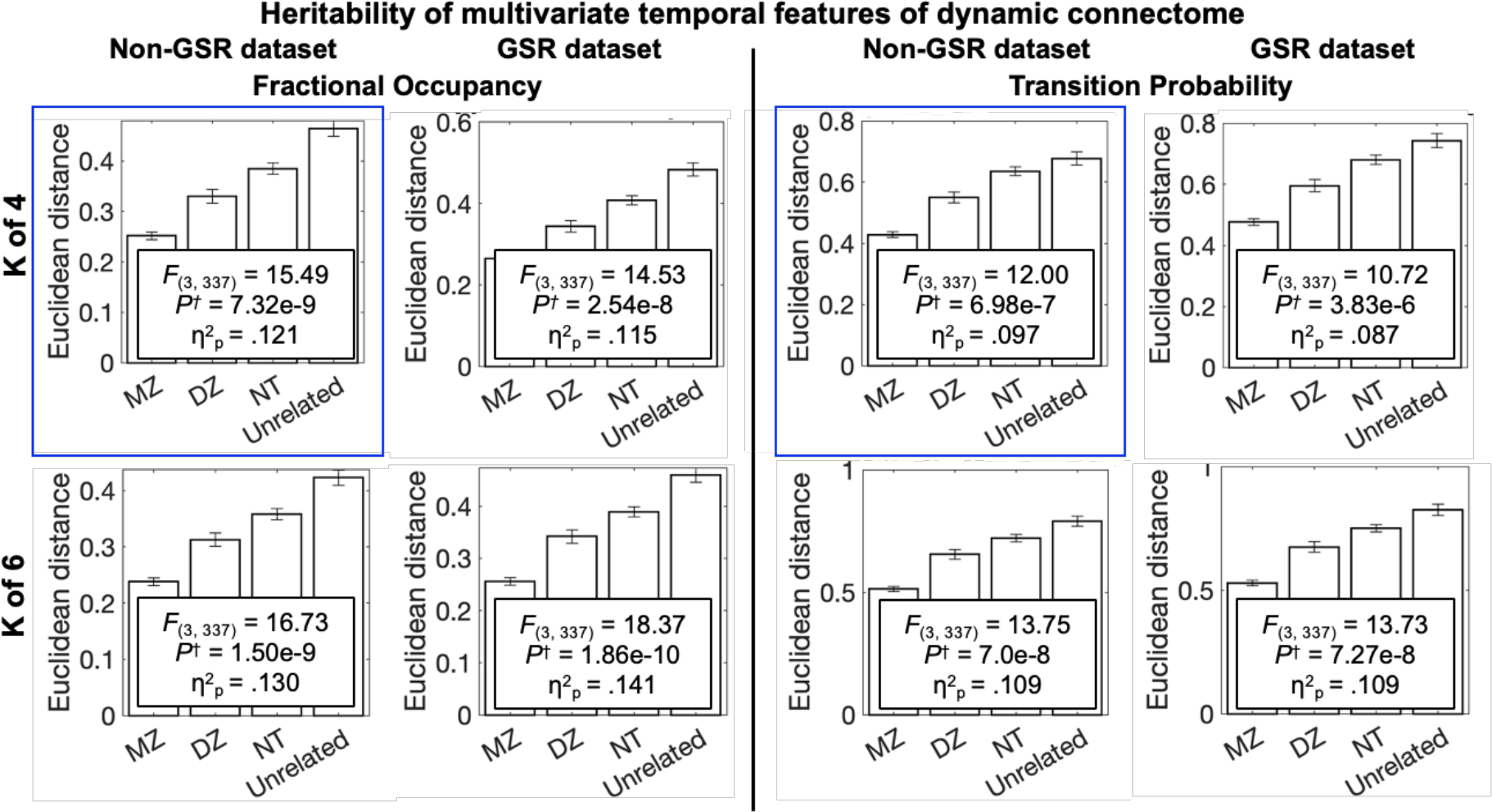
Heritability of temporal features of dynamic connectome. The bar plots show the similarity (= Euclidean distance) between subject pairs for fractional occupancy (FO) and transition probability (TP), respectively, across different combinations of the global signal regression (GSR) and the chosen number of discrete connectome states (K). Panels on the left of the vertical line are for FO, and those on the right are for TP. All panels robustly support that the strong heritability of the multivariate temporal features of dynamic connectome. *F* values are reported for one-way ANCOVAs of the factor sibling status (four levels: monozygotic twins (MZ), dizygotic twins (DZ), non-twin siblings (NT), and pairs of unrelated individuals), adjusted for age and head motion. K: the chosen number of discrete connectome states, GSR: global signal regression, *P*^†^: *P* values Bonferroni-corrected for four dependent variables (FO, TP, TV-FCDMN-FPN, and TV-Modularity). η^2^p: Partial Eta squared effect size. Note that the results highlighted in blue boxes are the same as reported in the main manuscript (Fig. 3) and provided again here for ease of comparison.

**Fig. S3.**
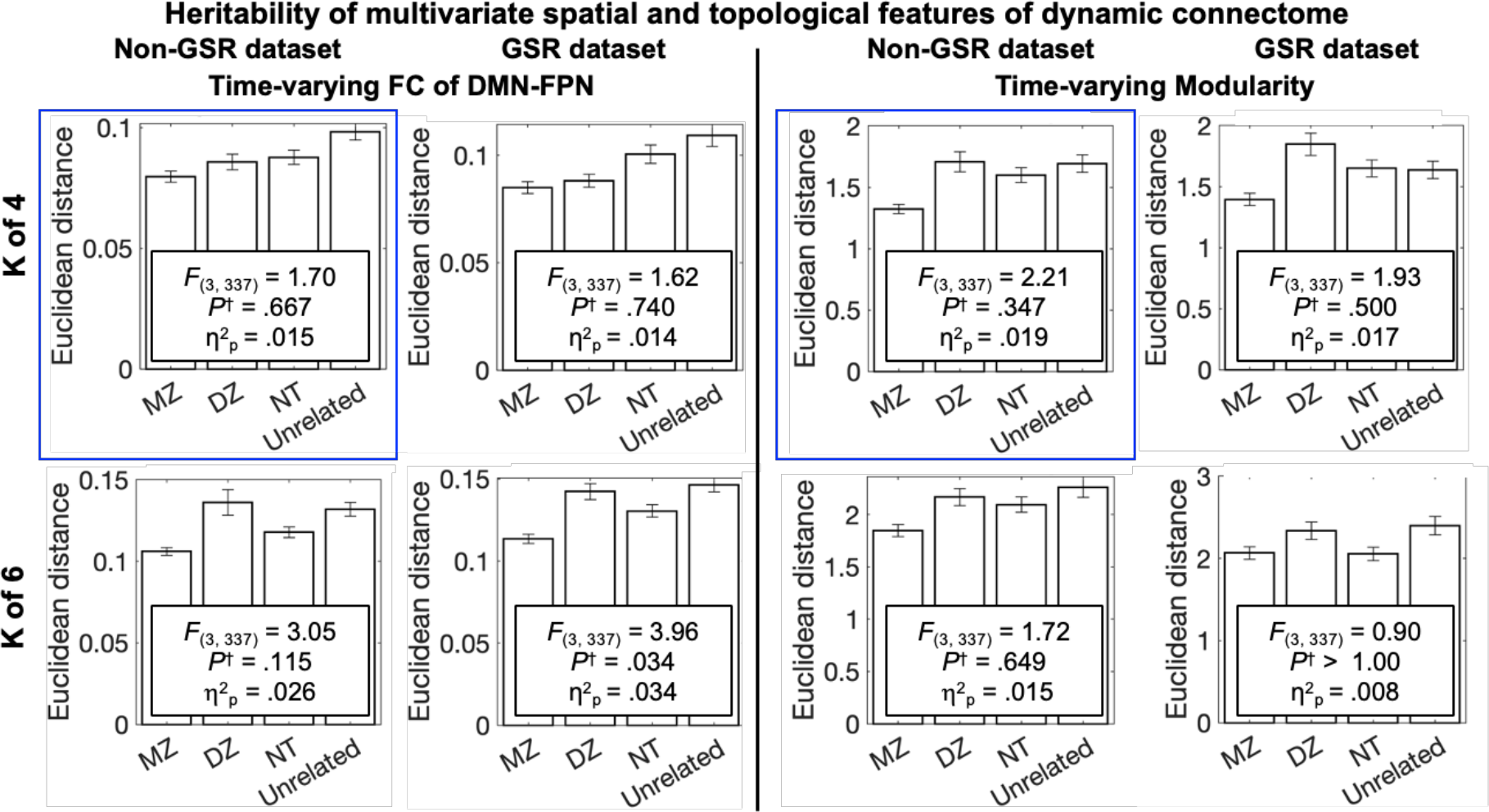
Heritability of spatial and topological features of dynamic connectome. The bar plots show the similarity (= Euclidean distance) between subject pairs for time-varying functional connectivity between default mode and frontoparietal networks (TV-FCDMN-FPN) and time-varying global modularity (TV-Modularity), respectively, across different combinations of the global signal regression (GSR) and the chosen number of discrete connectome states (K). Panels on the left of the vertical line are for the spatial feature, TV-FCDMN-FPN, and those on the right are for the topological feature, Modularity. Given the small effect size in the four-state models and the noticeably high similarity between dizygotic (DZ) twin pairs in the six-state models, we cautiously suggest that there is some evidence that the multivariate spatial feature is also heritable. Heritability of the global topological characteristic was not supported from our results. *F* values are reported for one-way ANCOVAs of the factor sibling status (four levels: monozygotic twins (MZ), dizygotic twins (DZ), non-twin siblings (NT), and pairs of unrelated individuals), adjusted for age and head motion. K: the chosen number of discrete connectome states, GSR: global signal regression, *P*^†^: *P* values Bonferroni-corrected for four dependent variables (FO, TP, TV-FCDMN-FPN, and TV- Modularity). η^2^p: Partial Eta squared effect size. Note that the results highlighted in blue boxes are the same as reported in the main manuscript (Fig. 3) and provided again here for ease of comparison.

**Fig. S4.**
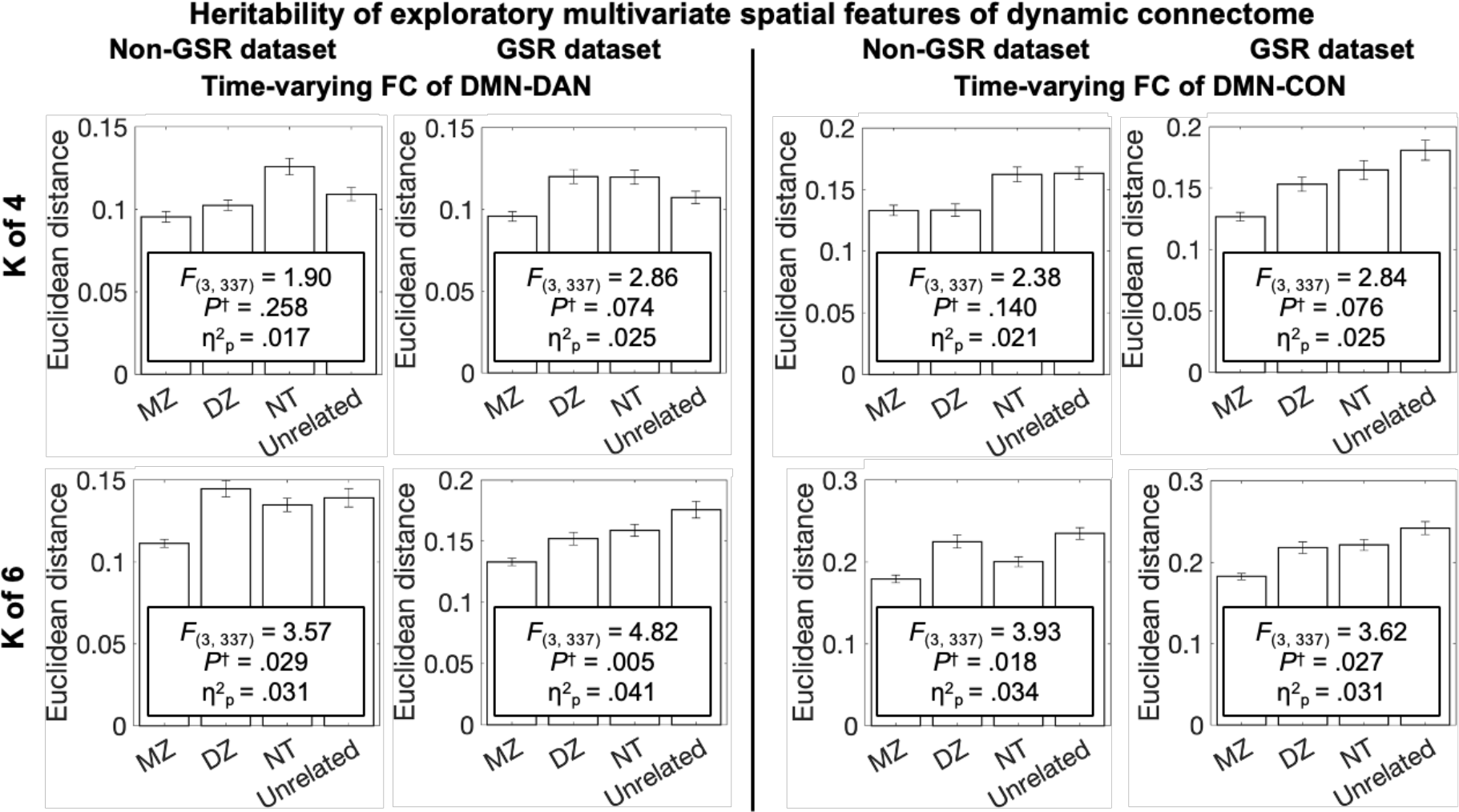
Exploring heritability of other spatial features: FCs between DMN and other top-down control networks. The bar plots show the similarity (= Euclidean distance) between subject pairs for time-varying functional connectivity between default mode and task-positive networks: dorsal attention network (TV-FCDMN-DAN) and cingulo-opercular network (TV-FCDMN-CON), respectively, across different combinations of the global signal regression (GSR) and the chosen number of discrete connectome states (K). Panels on the left of the vertical line are for TV-FCDMN-DAN, and those on the right are for TV-FCDMN-CON. Despite the small effect size (or no statistical significance) and the noticeably high similarity between dizygotic (DZ) twin pairs in some conditions, we cautiously suggest that there is some evidence that the multivariate spatial feature is also heritable. *F* values are reported for one-way ANCOVAs of the factor sibling status (four levels: monozygotic twins (MZ), dizygotic twins (DZ), non-twin siblings (NT), and pairs of unrelated individuals), adjusted for age and head motion. K: the chosen number of discrete connectome states, GSR: global signal regression, *P*^†^: *P* values Bonferroni-corrected for two dependent variables (TV-FCDMN-DAN, TV-FCDMN-CON. η^2^p: Partial Eta squared effect size.

**Fig. S5.**
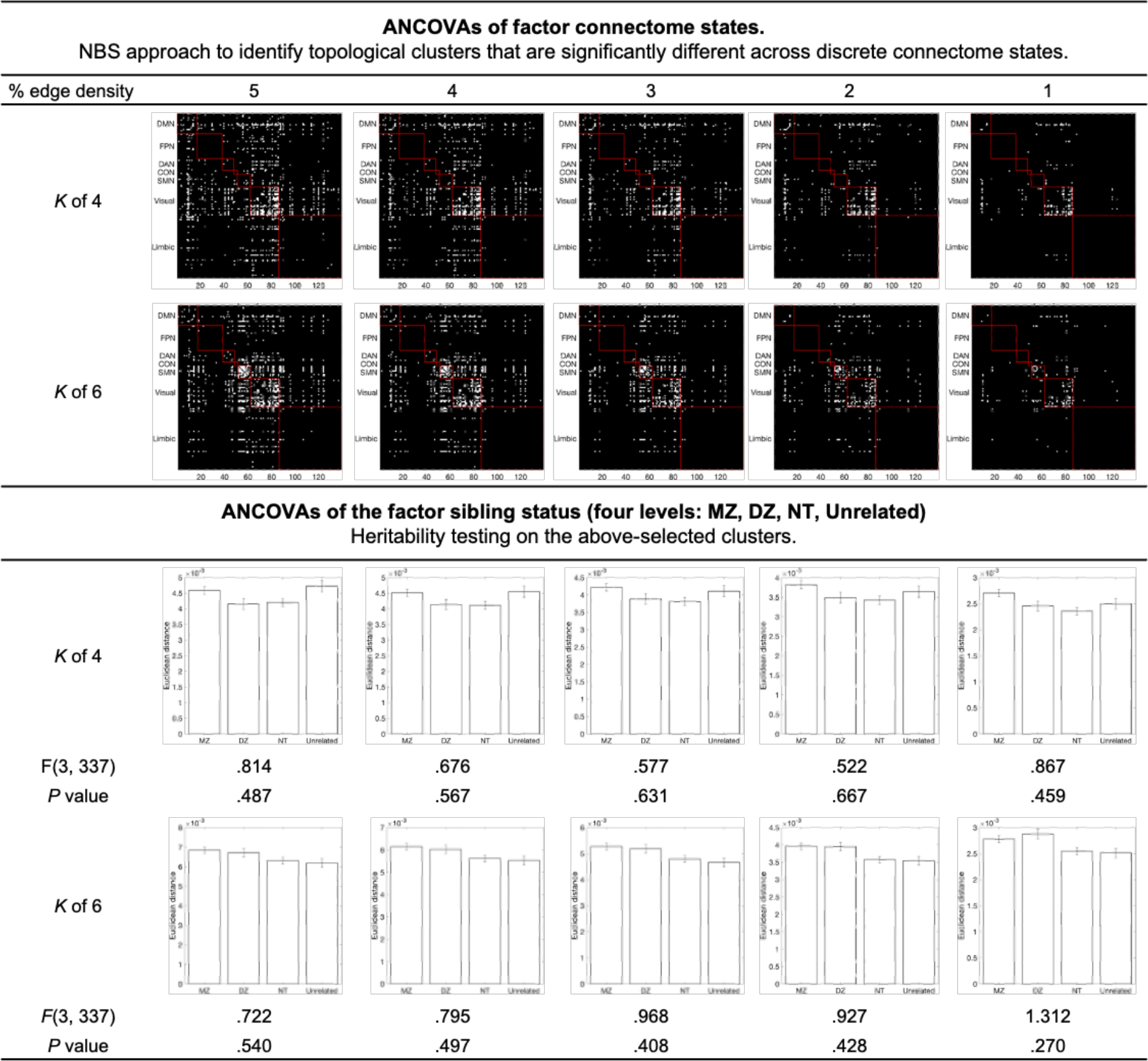
Exploring heritability of other spatial features: data-driven connectivity patterns. We performed network-based statistics (NBS) to select clusters, or connected sets of edges (region-pairs), in a data-driven manner for subsequent heritability analysis. We started with the absolute functional connectivity (FC) matrices, where all edges were transformed into absolute values to focus on the strength of connections, regardless of their connectivity direction. For each edge, an ANCOVA of the factor state was calculated to quantify the degree to which each edge’s connectivity strength differed across states, adjusted for age and head motion. The ensuing edgewise *F* statistics was threshold at arbitrary values, allowing 1 ∼ 5% of edges to survive. NBS was applied to the threshold matrices, resulting in one significant cluster at each threshold visualized as symmetric binary matrices. We averaged the absolute FC values of all edges in the cluster for each state, together constructing the multivariate spatial feature. The bar plots show the similarity (= Euclidean distance) of this multivariate feature between subject pairs. To assess heritability, *F* values are reported for one-way ANCOVAs of the factor sibling status (four levels: monozygotic twins (MZ), dizygotic twins (DZ), non-twin siblings (NT), and pairs of unrelated individuals). K: the chosen number of discrete connectome states.

**Fig. S6.**
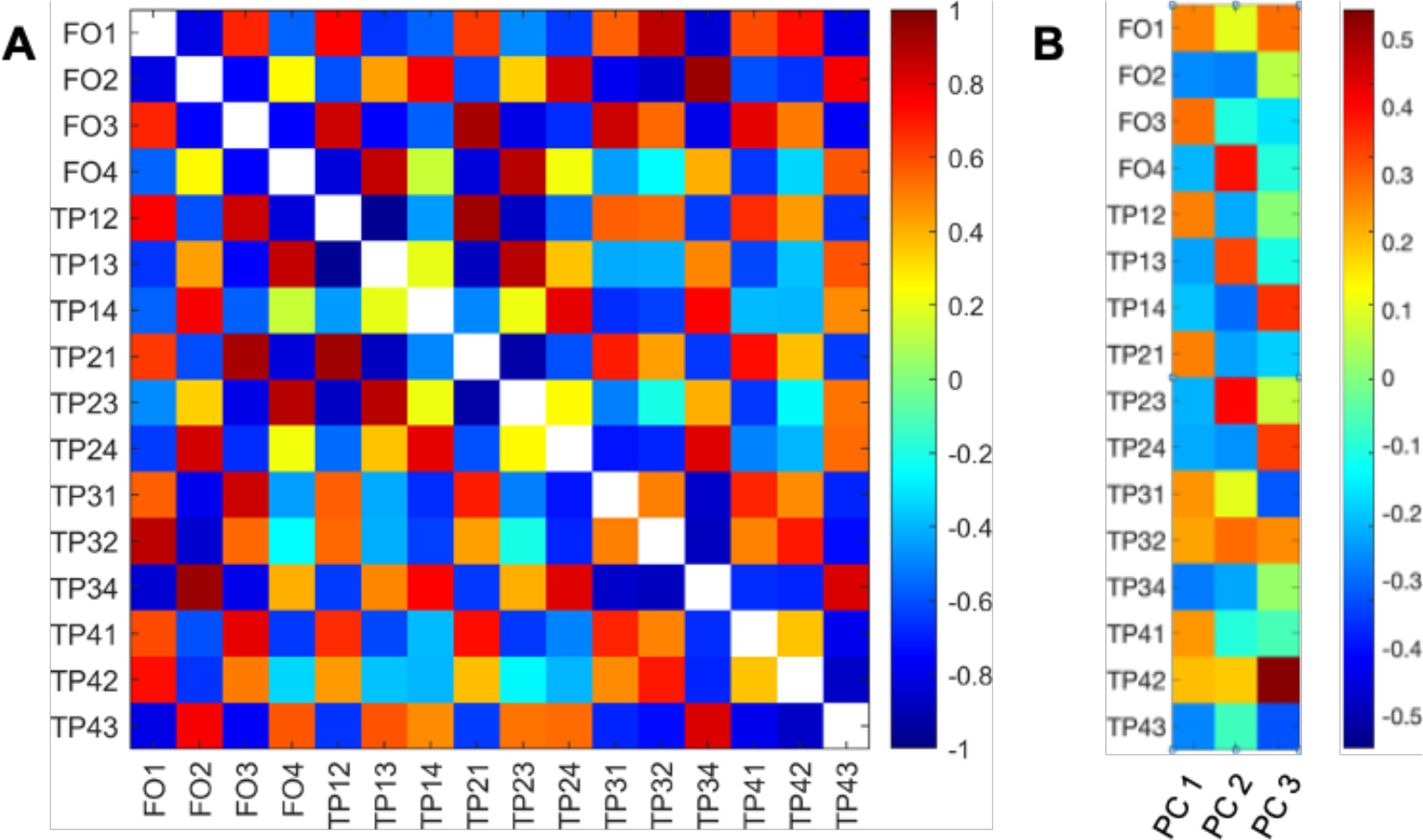
Overview of the temporal features of the dynamic connectome. (A) Pearson’s correlation matrix for the temporal features of the dynamic connectome obtained from the four-state model, color-coded for Pearson’s correlation coefficient (sample size *N* = 997). (B) Principal component (PC) coefficient matrix for the 3 PCs with PC variance > 1.

**Fig. S7.**
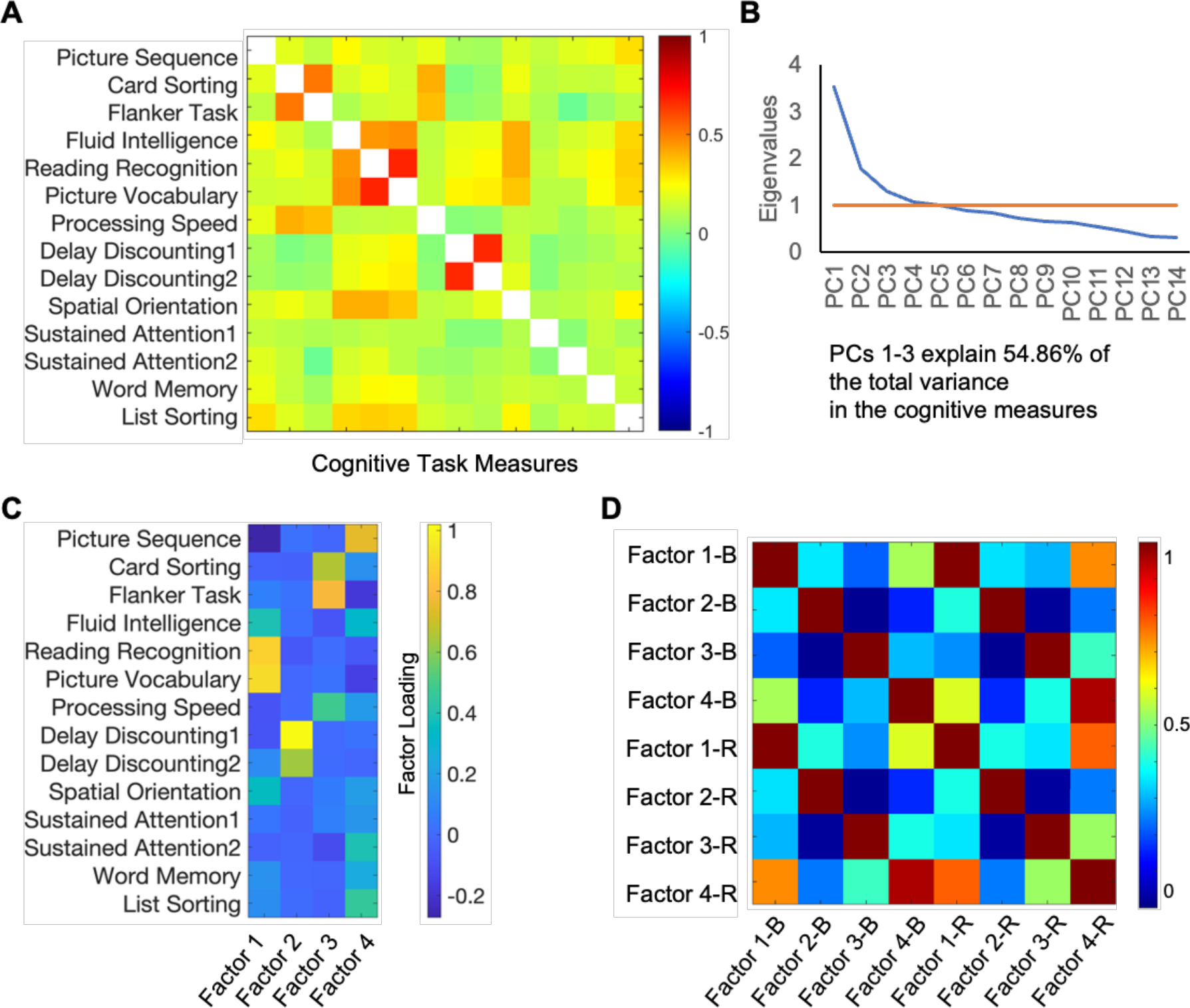
Overview of the cognitive dataset. (A) Empirical correlation matrix for the 14 cognitive performance measures in HCP (sample size *N* = 997), color-coded for Pearson’s correlation coefficient. (B) Scree plot of the eigenvalues of principal components obtained from the covariance matrix of the cognitive measures, where red colored reference line (y = 1) indicates the cut-off value used for the present study. (C) Factor loading matrix for 14 cognitive measures clustered into 4 factors based on ridge regression method. (D) Pearson’s correlation matrix for two sets of factor scores derived using Bartlett method (Factor 1-B to Factor 4-B) and ridge regression method (Factor1-R to Factor4-R). PC: principal component.

**Fig. S8.**
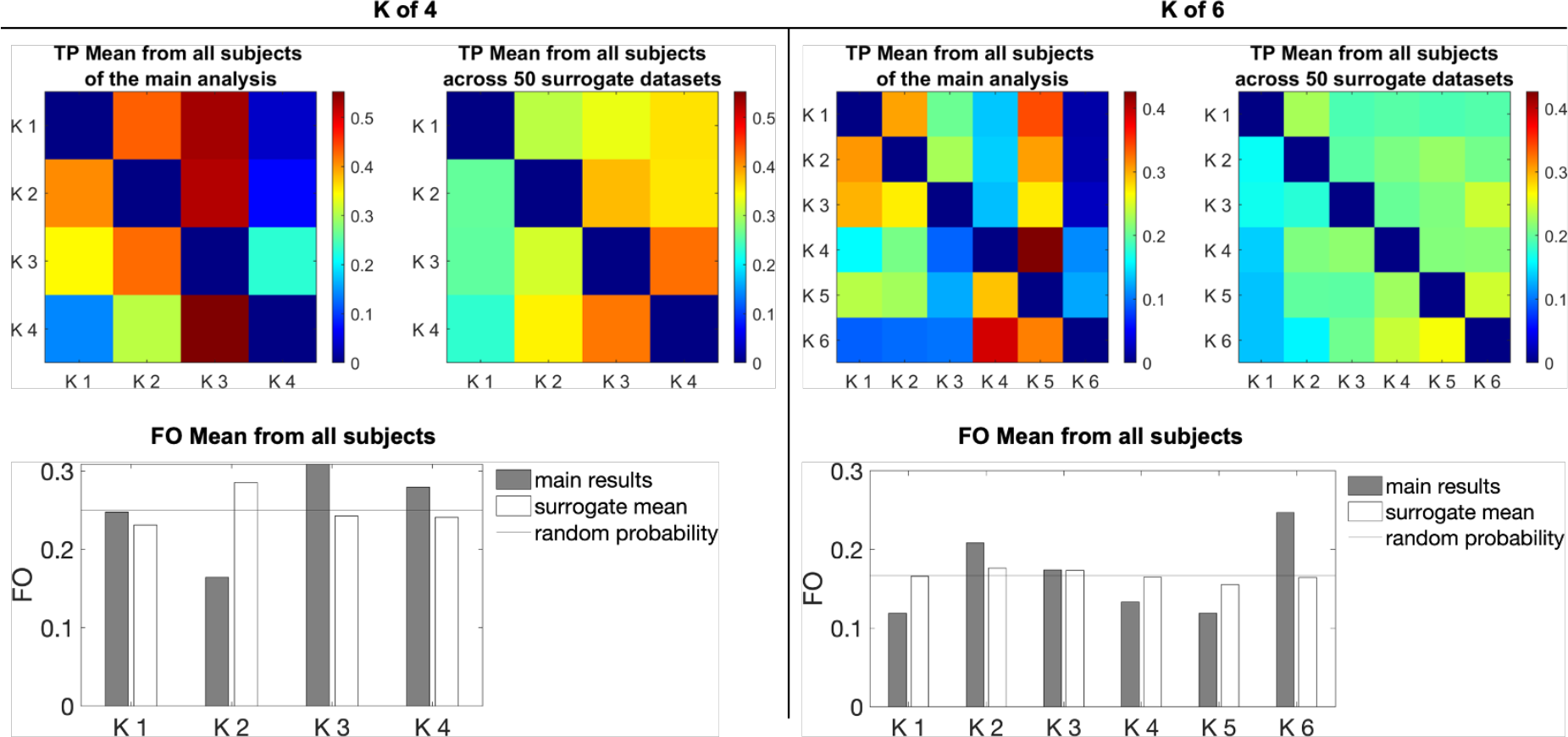
Distribution of temporal features of connectome dynamics from the null models. Difference between the TP matrix and FO distribution obtained from the non-GSR HCP dataset and those averaged from the 50 surrogate datasets. Transition probability matrix (TP) and fractional occupancy (FO) computed and averaged from the 50 surrogate data show probabilistic distribution, indicating the absence of meaningful dynamic properties in the surrogate data, whereas TP and FO obtained from the real data show non-random sequencing of brain networks.

## Supplementary Tables

**Table S1.**
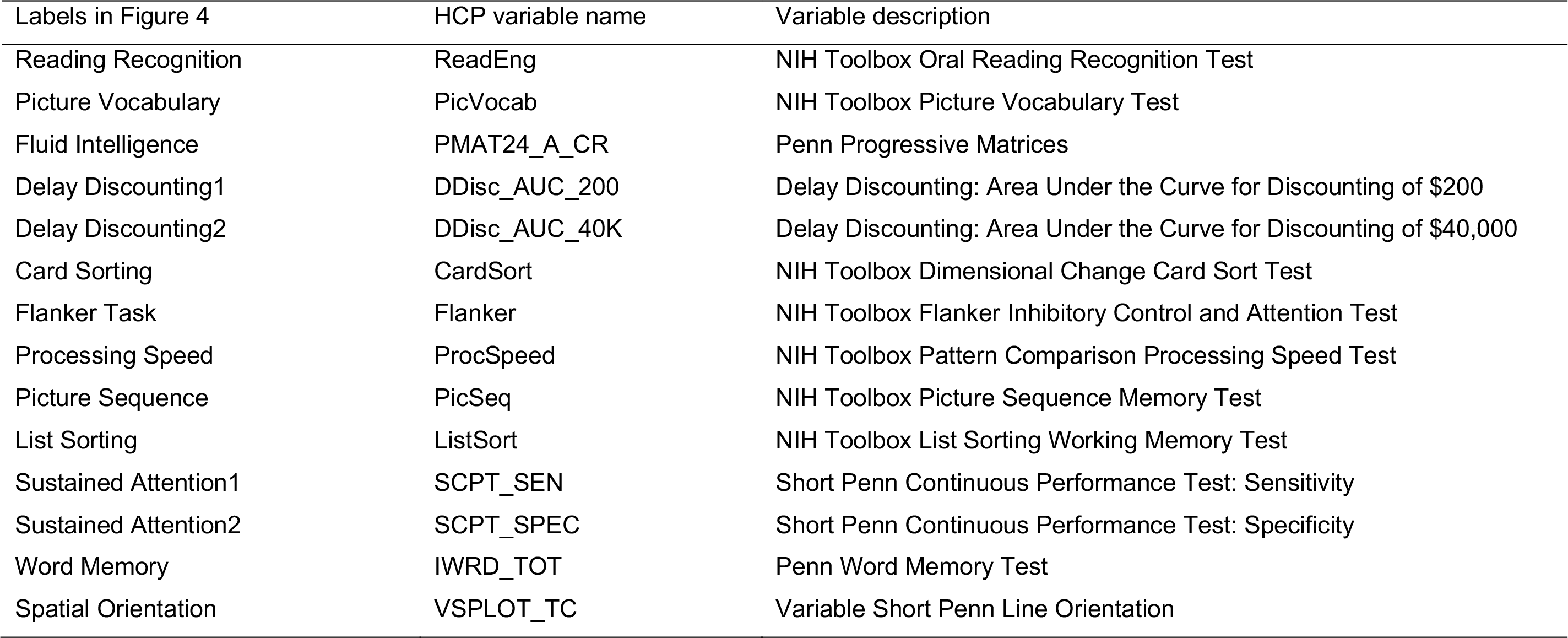
List of 14 cognitive measures from HCP and their basic descriptions.

**Table S2.** Bayes factor from Bayesian ANCOVAs of sibling status (four levels) reported for all investigated features (*K* = 4 stats, non-GSR data).

|  | Temporal features |  | Spatial features |  | Exploratory spatial features |  |  |  |  |  |  |
| --- | --- | --- | --- | --- | --- | --- | --- | --- | --- | --- | --- |
|  | FO | TP | TV-FC<sub>DMN-FPN</sub> | TV-Modularity | TV-FC<sub>DMN-CON</sub> | TV-FC<sub>DMN-DAN</sub> | NBS<br>5% | NBS<br>4% | NBS<br>3% | NBS<br>2% | NBS<br>1% |
| BF<sub>10</sub> | 1.96e+7 | 6.49e+6 | .12 | .47 | .47 | .85 | .04 | .03 | .03 | .03 | 0.27 |
| BF<sub>01</sub> | 5.11e-8 | 1.54e-7 | 8.042 | 2.146 | 2.134 | 1.183 | 25.23 | 29.91 | 32.36 | 31.10 | 19.23 |
| Error % | 2.18e-4 | .002 | .002 | 9.50e-6 | .062 | 3.01e-4 | .008 | .010 | .011 | .011 | .007 |

BF<sub>10</sub> is the probability of H<sub>1</sub> against H<sub>0</sub>, and BF<sub>01</sub> is the inverse i.e. the probability of H<sub>0</sub> against H<sub>1</sub>.
NBS denotes the significant cluster of edges extracted from Network Based Statistics (top x% of edges most dissimilar across states; cf. Fig. S5)

**Table S3.**
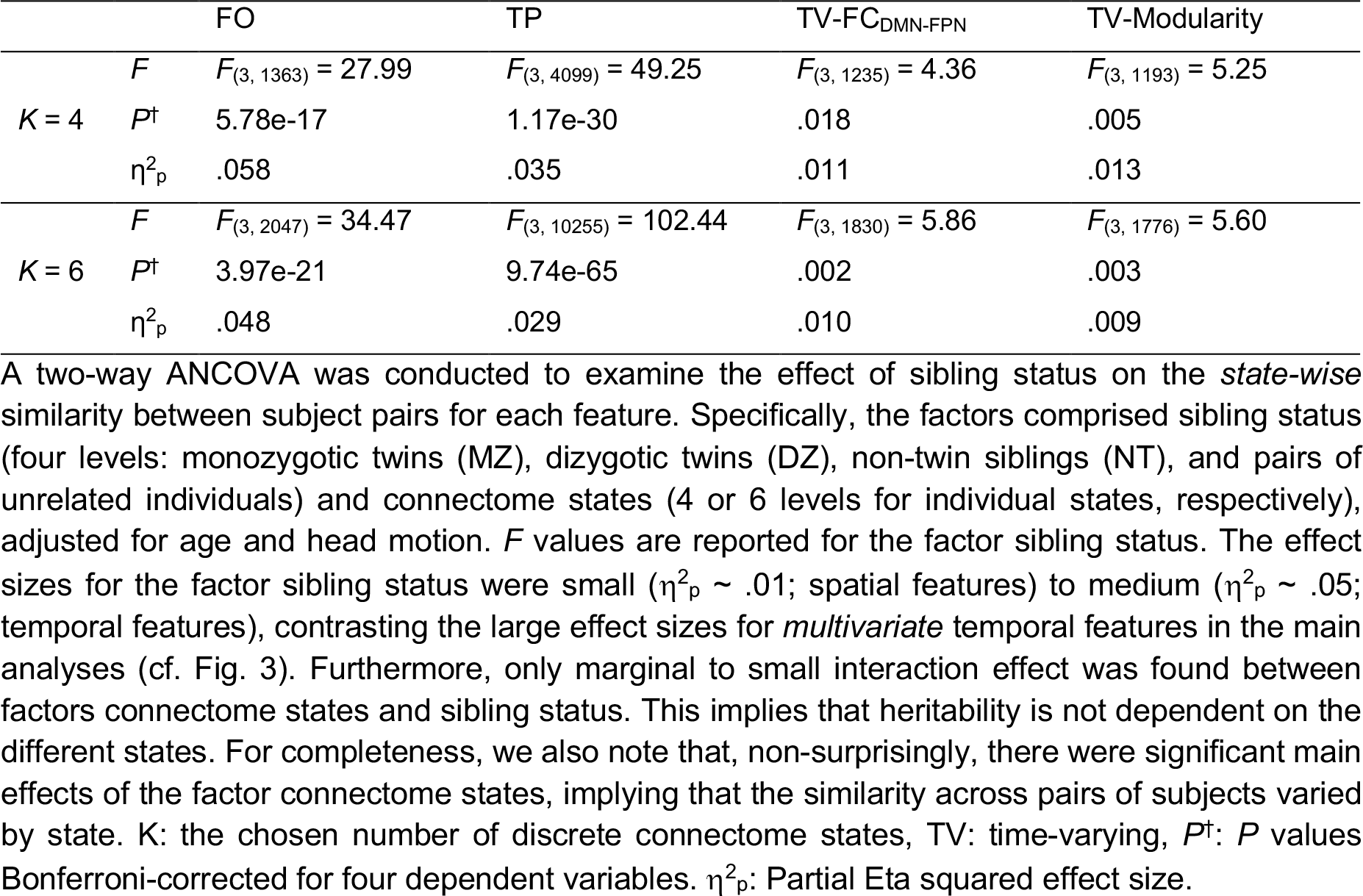
Heritability of individual components of dynamic connectome features.

**Table S4.** Cumulative weighted heritability of the individual components of dynamic connectome features in the four-state model.

|  | FO1 | FO2 | FO3 | FO4 |  |  |  |  |  |  |  |  |  |
| --- | --- | --- | --- | --- | --- | --- | --- | --- | --- | --- | --- | --- | --- |
| Model | ACE | ACE | ACE | ACE |  |  |  |  |  |  |  |  |  |
| $\sigma_m$ | .13 | .19 | .20 | .18 | | | | | | | | | |
| $\gamma_m$ | .18 | .27 | .29 | .26 | | | | | | | | | |
| $h^2$ | .640 | .377 | .347 | .592 | | | | | | | | | |
| CI | (.53, .75) | (.01, .74) | (-.06, .75) | (.48,.70) |  |  |  |  |  |  |  |  |  |
| weight*$h^2$ | .115 | .106 | .101 | .154 | | | | | | | | | |
| $\sum_m \gamma_m h_m^2$ | .476 | | | | | | | | | | | | |
|  | TP21 | TP31 | TP41 | TP12 | TP32 | TP42 | TP13 | TP23 | TP43 | TP14 | TP24 | TP34 |  |
| Model | ADE | ACE | ACE | ADE | ACE | ACE | ADE | ADE | ACE | ACE | ACE | ACE |  |
| $\sigma_m$ | .24 | .13 | .12 | .23 | .14 | .14 | .21 | .21 | .22 | .05 | .08 | .23 | |
| $\gamma_m$ | .12 | .07 | .06 | .11 | .07 | .07 | .10 | .10 | .11 | .02 | .04 | .11 | |
| $h^2$ or $H^2$ | .629 | .003 | .384 | .623 | .349 | .116 | .627 | .621 | .254 | .033 | .228 | .138 | |
| CI | (.53, .73) | (-.37, .38) | (-.07, .84) | (.52, .73) | (-.09, .79) | (-.37, .61) | (.53, .73) | (.52, .72) | (-.18, .69) | (-.46, .53) | (-.22, .67) | (-.25, .53) |  |
| weight*$H^2$ | .075 | .0002 | .023 | .069 | .024 | .008 | .063 | .062 | .028 | .0007 | .009 | .015 | |
| $\sum_m \gamma_m h_m^2$ | .377 | | | | | | | | | | | | |

$\sum_m \gamma_m h_m^2$ : the sum of weighted heritability of its individual components (Ge et al. 2016), where $\gamma_m = \frac{\sigma_m}{\sum_m \sigma_m}$ with $\sum_m \gamma_m = 1$ , and $h_m^2$ is the heritability of the $m^{\text{th}}$ state. *n.a.*, not applicable. $h^2$ : the narrow-sense heritability estimated as $\sigma^2_A/(\sigma^2_A + \sigma^2_c + \sigma^2_E)$ ; $H^2$ : the broad-sense heritability estimated as $(\sigma^2_A + \sigma^2_D)/(\sigma^2_A + \sigma^2_D + \sigma^2_E)$ ; CI: Confidence Interval (lower bound, upper bound).

